# Nuclear receptor NHR-49 promotes peroxisome proliferation to compensate for aldehyde dehydrogenase deficiency in *C. elegans*

**DOI:** 10.1101/2020.12.04.411637

**Authors:** Lidan Zeng, Xuesong Li, Christopher B. Preusch, Gary J. He, Ningyi Xu, Tom H. Cheung, Jianan Qu, Ho Yi Mak

## Abstract

The intracellular level of fatty aldehydes is tightly regulated to minimize the formation of toxic aldehyde adducts of cellular components. Accordingly, deficiency of a fatty aldehyde dehydrogenase FALDH causes the neurologic disorder Sjögren-Larsson syndrome (SLS) in humans. However, cellular responses to unresolved, elevated fatty aldehyde levels are poorly understood. Based on lipidomic and imaging analysis, we report that the loss of endoplasmic reticulum-, mitochondria- and peroxisomes-associated ALH-4, the *C. elegans* FALDH ortholog, increases fatty aldehyde levels and reduces fat storage. ALH-4 deficiency in the intestine, cell-nonautonomously induces NHR-49/NHR-79-dependent hypodermal peroxisome proliferation. This is accompanied by the upregulation of catalases and fatty acid catabolic enzymes, as indicated by RNA sequencing. Such a response is required to counteract ALH-4 deficiency since *alh-4; nhr-49* double mutant animals are not viable. Our work reveals unexpected inter-tissue communication of fatty aldehyde levels, and suggests pharmacological modulation of peroxisome proliferation as a therapeutic strategy for SLS.

## Introduction

Metabolic activities and environmental factors can cause oxidative stress through the production of reactive oxygen species (ROS). Although oxidative stress is often linked to cellular malfunction and organismal aging, it may also be integral to physiological processes such as stem cells maintenance and tissue differentiation (An and Blackwell, 2003; Bailey et al., 2015; Ushio-Fukai and Rehman, 2014). Therefore, deeply conserved signaling networks exist to monitor and resolve oxidative stress, in part through transcriptional outputs (Lehtinen et al., 2006; Rizki et al., 2012; Wang et al., 2013). For example, peroxisomal fatty acid β-oxidation produces hydrogen peroxide, which can be resolved through the action of catalases (Schrader and Fahimi, 2006; Wanders et al., 2015).

ROS can attack polyunsaturated fatty acids (PUFAs) that are associated with membrane lipids (Negre-Salvayre et al., 2008). Such peroxidation of PUFAs generates short chain and medium chain fatty aldehydes (Guéraud et al., 2010). In addition, long chain fatty aldehydes can be derived from plasmalogens (Ebenezer et al., 2020). Reactive fatty aldehydes form covalent adducts with proteins, lipids and nucleic acids and contribute to oxidative stress. Therefore, the level of cellular fatty aldehydes is under tight regulation. In humans, membrane-proximal fatty aldehydes can be neutralized by the fatty aldehyde dehydrogenase (FALDH), which belongs to the aldehyde dehydrogenase (ALDH) superfamily (Rizzo, 2014). ALDHs limit cellular stress by catalyzing the NAD(P)+ dependent conversion of fatty aldehydes to fatty acids, which can then be fed into anabolic or catabolic pathways (Singh et al., 2013). FALDH, also called ALDH3A2, is distinct from other human ALDHs because it associates with the ER or peroxisomal membranes (Ashibe et al., 2007; Costello et al., 2017). Besides ROS-induced aldehydes, FALDH processes fatty aldehydes from the metabolism of etherlipids (such as plasmalogens), sphingolipids and dietary branched-chained lipids (Rizzo, 2014). The importance of FALDH is underscored by the discovery that loss of function mutations in FALDH cause the Sjögren-Larsson syndrome in humans, who suffer from ichthyosis and neurologic disorder such as spasticity. FALDH is required for converting the sphingosine 1-phosphate metabolite hexadecenal to hexadecenoic acid in the conversion of sphingolipid to glycerolipid (Nakahara et al., 2012). Therefore, it has been proposed that the accumulation of hexadecenal due to FALDH deficiency contributes to the pathology of Sjögren-Larsson syndrome (James and Zoeller, 1997; Nakahara et al., 2012). However, the mechanisms by which hexadecenal harms the skin and nervous system are unclear. Furthermore, strategies to overcome FALDH deficiency have been lacking.

Nuclear receptors are conserved metazoan transcription factors that are distinguished by their C2H2 zinc finger-containing DNA binding domain and ligand binding domain (Mangelsdorf et al., 1995). In humans, nuclear receptors, such as estrogen receptors, are known to bind steroid hormones with high affinity. This is because of the tight fit between the ligand binding pocket with the cognate ligand. However, there were also a number of nuclear receptors, called orphan receptors, which had no known ligands when they were initially identified (Evans and Mangelsdorf, 2014). Subsequently efforts led to the discovery of fatty acid or cholesterol metabolites as previously unrecognized ligands to some of them. Prime examples include the peroxisome proliferator activated receptors (PPARs). The structures of the ligand binding domain of PPARs reveal a large ligand binding pocket that accommodates a range of fatty acid derivatives and synthetic ligands (Xu et al., 2001). PPARs form heterodimers with retinoid X receptors (RXRs) and bind to PPAR response elements (PPREs) to modulate the expression of their target genes (Kliewer et al., 1992; Kojetin et al., 2015). Based on genetic and pharmacological studies in vitro, in mice and humans, PPARα and PPARγ have well-established roles for energy homeostasis (Dubois et al., 2017; Jeninga et al., 2009). For example, PPARα is pivotal for the activation of genes that are required for fatty acid uptake and turnover (Pawlak et al., 2015).

The nuclear receptor family is greatly expanded in *C. elegans*. In comparison to 48 nuclear receptors in humans, the *C. elegans* genome is predicted to encode 284 nuclear receptors (Taubert et al., 2011). To date, most of them remain uncharacterized and have no known ligands. Nevertheless, a handful of *C. elegans* nuclear receptors have been shown to regulate developmental dauer arrest, molting, aging or metabolism (Brock et al., 2006; Folick et al., 2015; Hayes et al., 2006; Liang et al., 2010; Lin and Wang, 2017; Magner et al., 2013; Motola et al., 2006; Mullaney et al., 2010). Thus, *C. elegans* nuclear receptors are pervasive in facilitating inter-tissue communication, similar to their mammalian counterparts.

One of the best characterized *C. elegans* nuclear receptors is NHR-49. Although it bears relatively high sequence similarity to mammalian Hepatocyte Nuclear Factor 4 (HNF4), NHR-49 appears to serve similar functions as PPARα (Van Gilst et al., 2005a). Notably, NHR-49 modulates gene expression programs that govern fatty acid catabolism, desaturation and remodeling under fed and fasted conditions (Pathare et al., 2012; Ratnappan et al., 2014; Van Gilst et al., 2005b, 2005a). As a result, NHR-49 has been implicated in cold adaptation of phospholipid membrane composition and preservation of germline stem cells in starved animals (Angelo and Van Gilst, 2009; Svensk et al., 2013). NHR-49 has also been shown to regulate lifespan in response to diffusible molecules (Burkewitz et al., 2015; Qi et al., 2017). NHR-49 partners with different nuclear receptors to regulate distinct sets of target genes (Pathare et al., 2012). Here, we report that NHR-49 acts with NHR-79 to promote peroxisome proliferation in mutant worms that lack ALH-4, which is the predicted ortholog of mammalian FALDH. Surprisingly, ALH-4 deficient worms were less sensitive to specific oxidative stress than wild type worms, which could be attributed in part to NHR-49 dependent peroxisome proliferation. Accordingly, we found that *alh-4; nhr-49* mutant worms were genetically lethal. Our results reveal a nuclear receptor coordinated peroxisome proliferative response, which effectively counters the accumulation of toxic fatty aldehydes.

## Results

### Molecular cloning of *alh-4*

The attenuation of peroxisomal β-oxidation causes excess accumulation of TAG in expanded LDs in mutant animals that lack the DAF-22 thiolase (Zhang et al., 2010). This phenotype is most pronounced in the *C. elegans* intestine, which serves as a major fat storage tissue, and could be readily detected using quantitative Stimulated Raman Scattering (SRS) microscopy (Li et al., 2015) (Fig 1A). To identify genetic regulators of fat storage and LD expansion, we conducted a forward genetic screen for suppressor mutants that reversed the *daf-22* mutant phenotypes. A complementation group with three recessive alleles (*hj27*, *hj28*, *hj29*) were identified. Molecular cloning revealed that *hj27* is a missense allele and *hj28* and *hj29* are nonsense alleles of *alh-4* (Fig S1A), which encodes a predicted aldehyde dehydrogenase that catalyzes fatty aldehyde into fatty acid. Using CRISPR-mediated genome editing (Dickinson et al., 2013), we generated an additional *alh-4* deletion allele, *hj221* (Fig S1A). We found that *alh-4(hj221)*, as well as RNA interference (RNAi) against *alh-4*, suppressed LD expansion of *daf-22(−)* mutant animals (Fig S1B). Therefore, *alh-4(hj29)* is likely to be a strong loss of function allele.

**Figure 1.**
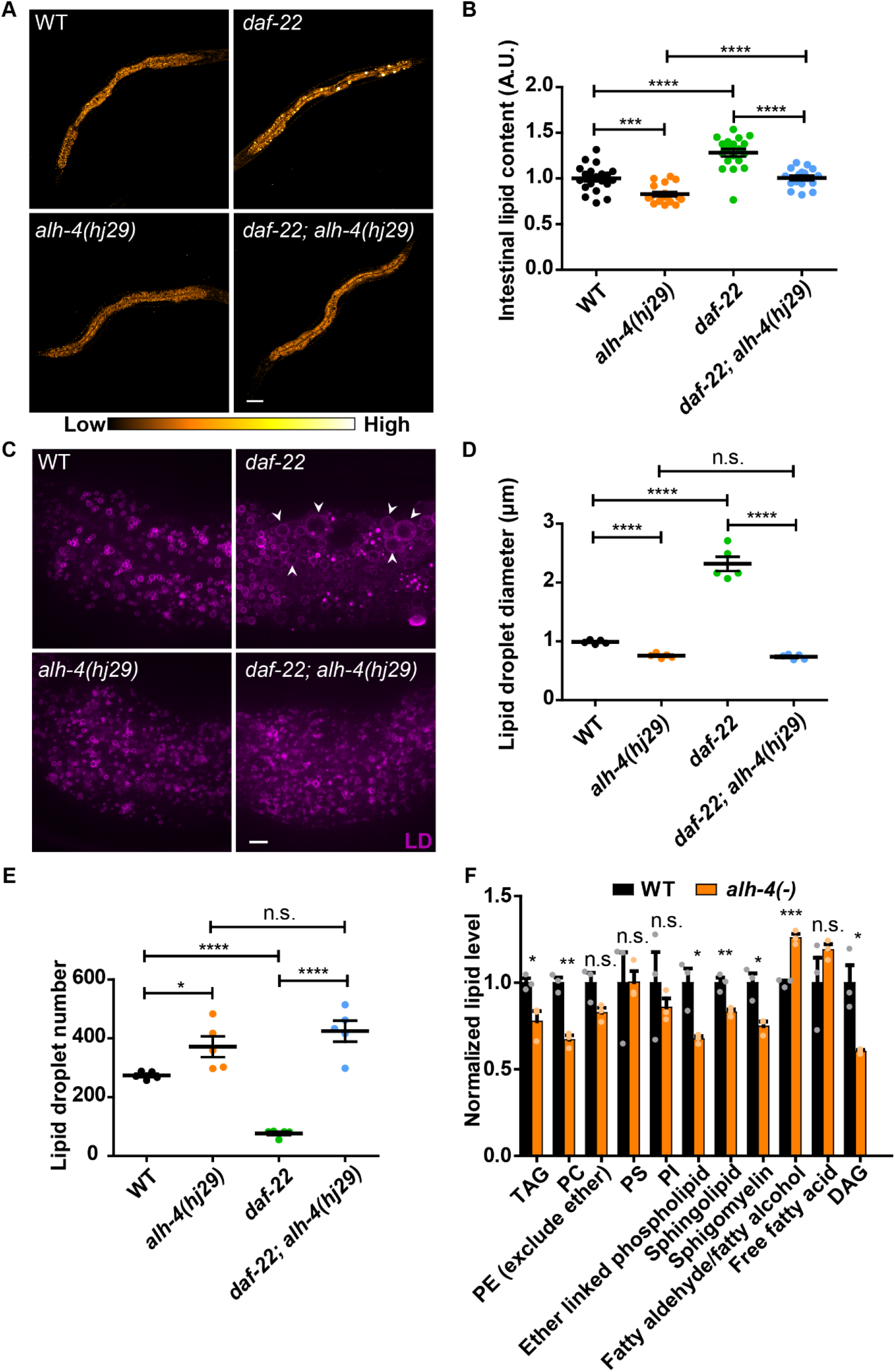
ALH-4 regulates intestinal lipid storage. (A) Representative Stimulated Raman Scattering (SRS) images of L4 worms of wild type (WT), *daf-22(ok693), alh-4*(*hj29)* and *alh-4(hj29); daf-22(ok693)*. Scale bar = 50μm. (B) Quantification of average SRS signal intensity. n = 20 for all groups. (C) Representative images of lipid droplets in the second intestinal segment of L4 worms. Lipid droplets (LDs) are labeled with mRuby::DGAT-2*(hjSi112)*. Enlarged LDs are marked by arrowheads. Scale bar = 5μm. (D) Quantification of lipid droplet diameter in the second intestinal segment. n = 5 for all groups. (E) Quantification of total lipid droplet number in the second intestinal segment in individual worms. n = 5 for all groups. (F) Lipidomic analysis of WT and *alh-4(−)* worms. Quantification of lipid species detected by LC-MS. The value of each species in wild type samples was set as 1. Each dot represents the average value of two technical replicates of each biological sample. n = 3 for all groups. TAG, triacylglycerol; PE, phosphatidylethanolamine; PC, phosphatidylcholine; PS, phosphatidylserine; PI, phosphoinositide; DAG, diacylglycerol. In all plots, mean ± SEM of each group is shown. For all statistical analysis, *p < 0.05, **p < 0.01, ***p < 0.001; ****p < 0.0001, n.s. not significant (unpaired Student’s t-test).

The loss of *alh-4* function increased the brood size and the rate of larval development of *daf-22(−)* worms (Fig S1D-E). Using SRS microscopy, we quantitatively measured the effect of ALH-4 deficiency on cellular fat storage. The loss of *alh-4* function suppressed fat accumulation and LD expansion of *daf-22(−)* worms (Fig 1A-D). In addition, the *alh-4(hj29)* worms accumulated less fat than wild type worms (Fig 1A-B). Accordingly, *alh-4(hj221)* (denoted as *alh-4(−)* hereafter) worms had significantly less TAG than wild type worms, based on liquid chromatography-mass spectrometry (LC-MS) analysis (Fig 1F). Since there was no significant difference in the pharyngeal pumping rate of wild type and *alh-4* mutant worms (Fig S1F), we concluded that the reduction in fat storage was unlikely due to a change in food intake. To compare the size of intestinal LDs in wild type and mutant worms, we used an established LD marker, mRuby::DGAT-2 (Klemm et al., 2013). In agreement with our observations with SRS, ALH-4 deficiency reduced the average intestinal LD size of wild type and *daf-22* mutant worms (Fig 1C-D). The LD size profiles of *alh-4(hj29)* and *daf-22*(−); *alh-4(hj29)* worms almost completely overlapped (Fig S1C). The reduction of LD size was accompanied by an increase in LD number in *alh-4(hj29)* and *daf-22*(−); *alh-4(hj29)* worms, in comparison with wild type and *daf-22*(−) worms, respectively (Fig 1E). We concluded that the loss of *alh-4* function suppressed LD expansion without compromising LD biogenesis.

### Tissue specific localization of ALH-4

Based on sequence homology, ALH-4 appears to be closely related to human Class III aldehyde dehydrogenases. In agreement with such prediction, LC-MS based lipidomic analysis indicated that *alh-4(−)* worms accumulated significantly more fatty aldehyde and its related metabolite fatty alcohol, than wild type worms (Fig 1F, S1G). Furthermore, ALH-4 and human ALDH3A2 (also known as FALDH) are the only proteins in this class of enzymes that bear C-terminal tail anchors in the form of a single transmembrane alpha helix. Prior studies indicated that ALDH3A2 associates with peroxisomal membrane and the endoplasmic reticulum via distinct C-terminal tails that are encoded by alternatively spliced exons (Ashibe et al., 2007). Similarly, we noted that three *alh-4* isoforms were annotated, based on publicly available RNA sequencing data (Fig 2A). These isoforms differ by their 3’ ends, which encode alternative C-termini that follow a common transmembrane helix. As a first step of determining the expression pattern and intracellular localization of ALH-4 isoforms, we inserted the coding sequence of the green fluorescent protein (GFP) immediately after the start codon of *alh-4* by CRISPR-mediated genome editing (Fig 2A). As a result, all three isoforms of ALH-4 were expressed as GFP fusion proteins from the endogenous locus. This GFP knockin strain, *alh-4(hj179)*, retained normal ALH-4 function because lipid droplet expansion was unperturbed when *daf-22* was mutated, unlike what we observed in *alh-4(−)* worms. We detected GFP::ALH-4 expression in the intestine and hypodermis, but not in the germline (Fig 2B-F). Our results suggest that ALH-4 may be important in regulating somatic lipid metabolism.

**Figure 2.**
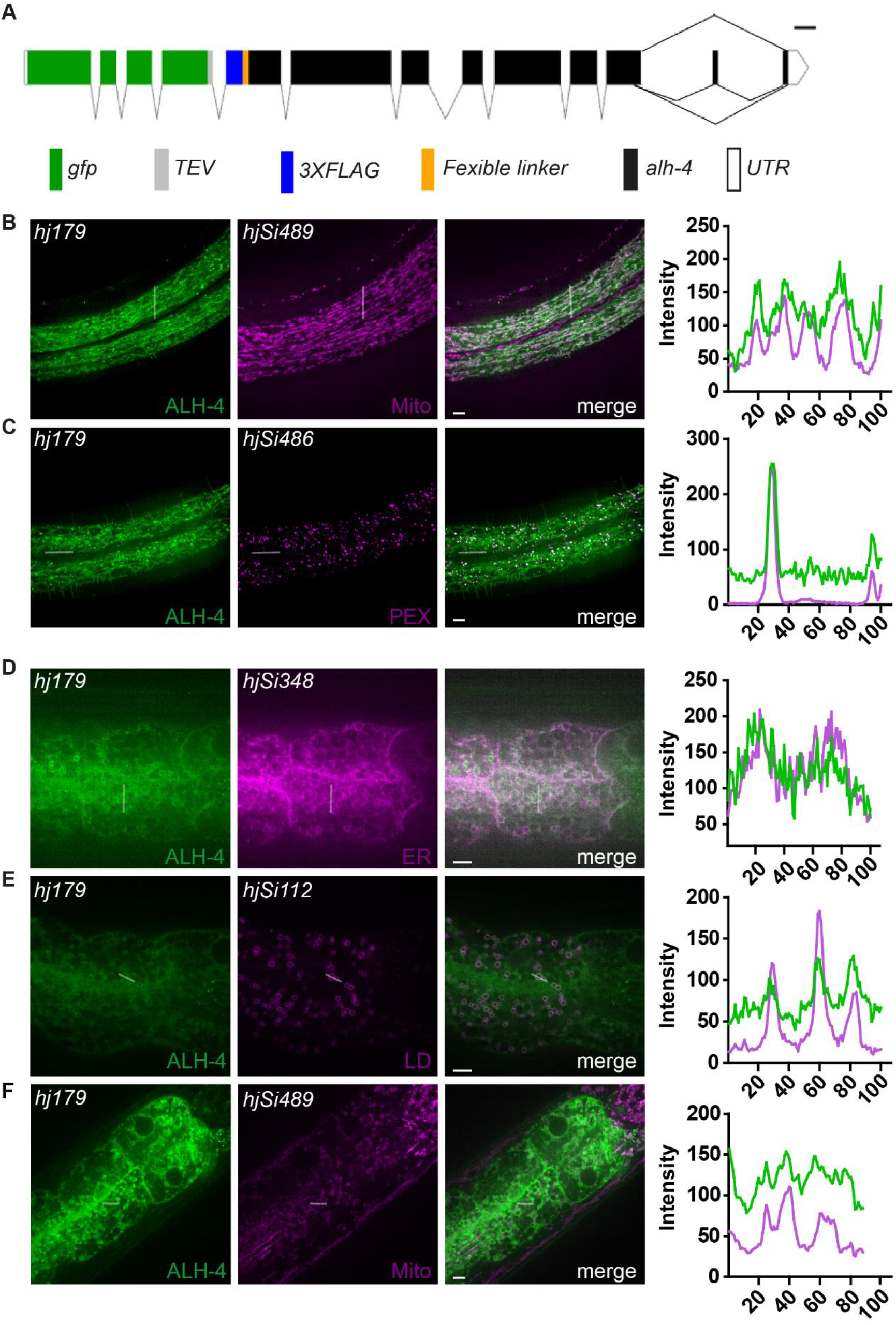
Tissue-specific subcellular localization of ALH-4. (A) Schematic representation of *gfp::alh-4(hj179)*. Scale bar = 100bp. Alternative splicing results in three ALH-4 isoforms that differ only at the C-terminus. (B-C) Representative images showing the subcellular localization of GFP::ALH-4 in the hypodermis (left). Fluorescence intensity profile from the indicated line scans (right). (B) GFP::ALH-4 largely colocalized with the mitochondria marker, TOMM-20N::mRuby (*hjSi489*). (C) The bight puncta of GFP::ALH-4 colocalized with the peroxisome marker, tagRFP::PTS1 (*hjSi486*). (D-F) Representative images showing the subcellular localization of GFP::ALH-4 in the intestine (left). Fluorescence intensity profile from the indicated line scans (right). (D) GFP::ALH-4 largely colocalized with the ER marker, ACS-22::tagRFP *(hjSi348)*. (E) Partial colocalization of GFP::ALH-4 with the lipid droplet marker mRuby::DGAT-2 (*hjSi112*). (F) GFP::ALH-4 did not colocalize with the mitochondria marker, TOMM-20N::mRuby (*hjSi489*). Scale bar = 5μm. Mito, mitochondria; PEX, peroxisome; ER, endoplasmic reticulum; LD, lipid droplet.

We used a suite of red fluorescent organelle markers to determine the subcellular localization of GFP::ALH-4. Specifically, mRuby::DGAT-2, ACS-22::tagRFP, and TOMM-20(N)::mRuby, tagRFP::peroxisomal targeting signal 1(PTS1) were used to label LDs, ER, mitochondria and peroxisomes, respectively (Cao et al., 2019; Xu et al., 2012). Live transgenic worms were observed using confocal microscopy. In the hypodermis, we detected co-localization of GFP::ALH-4 signals with red fluorescent signals that labeled mitochondria and peroxisomes (Fig 2B-C). In the intestine, GFP::ALH-4 signals co-localized with the ER marker and to a lesser extent, LD and mitochondrial markers (Fig 2D-F). Therefore, ALH-4 appeared to associate with an overlapping set of organelles, where lipid anabolism, storage or catabolism take place. It is conceivable that the fatty acid products of ALH-4 can be channeled into metabolic pathways locally at respective organelles.

### Determinants of organelle specific targeting of ALH-4 isoforms

Next, we determined if each of the three alternative C-termini of ALH-4 was responsible for its targeting to distinct organelles. To this end, we generated single-copy transgenes that placed the expression of GFP::ALH-4A, GFP::ALH-4B or GFP:ALH-3C under the control of the ubiquitous *dpy-30* promoter (Fig S2A). As a result, we were able to observe the localization of each isoform in the intestine and hypodermis at the same time. We found ALH-4A at mitochondria and peroxisomes in both tissues (Fig S2B-E). In contrast, ALH-4B was found primarily at the ER in the intestine (Fig S2F). However, we could not observe the association of ALH-4B with specific organelles in the hypodermis (Fig S2G). Finally, ALH-4C showed tissue specific targeting. In the intestine, ALH-4C was found at mitochondria and LDs (Fig S2H-I). However, it was only found at mitochondria in the hypodermis (Fig S2J). As a complementary approach, we generated three additional single-copy transgenes that drove the ubiquitous expression of GFP fused to the ALH-4 transmembrane helix and one of the three alternative C-termini (Fig S2L). In both intestine and hypodermis, we found that the C-terminus of ALH-4B was sufficient to target GFP to the ER (Fig S2M-N). In contrast to the full length protein, the C-terminus of ALH-4A could only target GFP to mitochondria, while the C-terminus of ALH-4C to mitochondria and the ER. Therefore, the targeting of full-length ALH-4A to peroxisomes, and ALH-4C to LDs appeared to require additional features beyond their C-termini. These features may be important for steering the respective isoforms directly to the second location. Alternatively, they may be required for the transfer of mitochondrial ALH-4 to peroxisomes or LDs via yet to be defined contact sites. Taken together, we conclude that the C-terminus of each ALH-4 isoforms serves as a tail anchor to membranes of distinct organelles.

### Membrane anchoring is required for ALH-4 function

ALH-4 is the only aldehyde dehydrogenase in *C. elegans* that is predicted to harbor a transmembrane helix. Its broad association with membranous organelles led us to ask if membrane association was required for ALH-4 function. Starting with the GFP knockin strain, *alh-4(hj179)*, which expressed full-length GFP::ALH-4 fusion protein from the endogenous locus, we used CRISPR to delete the sequence encoding the transmembrane helix and C-termini of the *alh-4* gene (Fig S3A). The GFP::ALH-4(ΔC) truncated protein retained the entire catalytic domain of ALH-4. As expected, the GFP fusion protein was cytosolic in both intestine and hypodermis (Fig 3A). The expression level of GFP::ALH-4(ΔC) was comparable to GFP::ALH-4 based on the intensity of GFP fluorescence. To assess the function of GFP::ALH-4(ΔC), we used mRuby::DGAT-2 as the LD marker. The average LD size was significantly reduced in animals that expressed GFP::ALH-4(ΔC), in comparison to those that expressed wild type GFP::ALH-4 (Fig 3B-C). The mutant phenotype was not as pronounced as in *alh-4(hj29)* worms, suggesting that the function of GFP::ALH-4(ΔC) was severely compromised but not eliminated.

**Figure 3.**
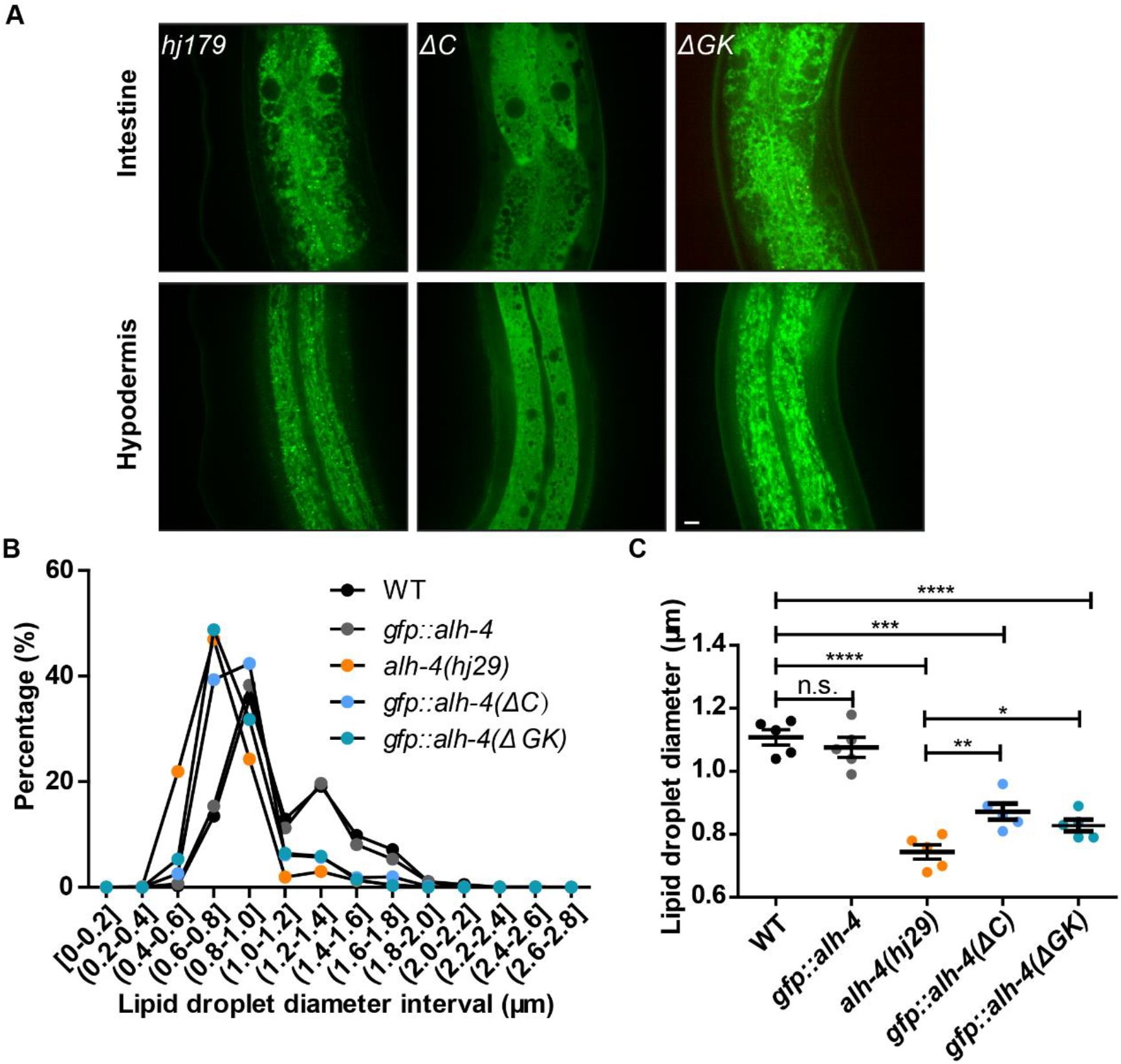
Membrane anchor and gatekeeper regions are necessary for proper ALH-4 function. (A) Representative images showing the subcellular localization of GFP tagged wild type ALH-4 (*hj179*) (left), cytosolic ALH-4 harboring a C-terminal deletion (ALH-4ΔC, *hj261*) (middle) and ALH-4 with a deletion of the gatekeeper region (ALH-4ΔGK, *hj264*) (right). Organelle-specific targeting was not affected by the deletion of the gatekeeper region. (A) The size distribution of lipid droplets in the second intestinal segment. mRuby::DGAT-2 (*hjSi112*) was used to label lipid droplets. Scale bar = 5μm. (C) Quantification of lipid droplet size in the second intestinal segment. Both *gfp::alh-4(ΔC)* and *gfp::alh-4(ΔGK)* showed reduced lipid droplet size as observed in *alh-4*(*hj29*). *gfp::alh-4(hj179)* showed no significant difference in lipid droplet size comparing to WT, indicating that the function of ALH-4 was not compromised by GFP tagging. In the plot, mean ± SEM of each group is shown. For all statistical analysis, *p < 0.05, **p < 0.01, ***p < 0.001; ****p < 0.0001, n.s. not significant (unpaired Student’s t-test).

Structure-function analysis of ALDH3A2 indicated that a ‘gate-keeper’ helix is responsible for substrate selectivity in vitro (Keller et al., 2014). The helix is located immediately N-terminal to the transmembrane helix, and appears to be conserved in ALH-4 (Fig S3B). Therefore, we tested if the gate-keeper helix was required for ALH-4 function in vivo, by deleting its coding sequence from the GFP knockin strain, *alh-4(hj179)*. We found that the organelle targeting of GFP::ALH-4(ΔGK) was unperturbed in intestine and hypodermis (Fig 3A). In addition, the expression level of GFP::ALH-4(ΔGK) was comparable to GFP::ALH-4 based on the intensity of GFP fluorescence. However, worms expressing GFP::ALH-4(ΔGK) had significantly smaller LDs when compared with those expressing GFP::ALH-4 (Fig 3B-C). Taken together, our results indicate that membrane anchoring by the transmembrane helix, and the gate-keeper helix are both required for ALH-4 function in vivo.

### Upregulation of a peroxisome-related gene expression program in ALH-4 deficient worms

The membrane anchored aldehyde dehydrogenase family, to which ALH-4 belongs, is known to play a critical role in converting membrane associated long-chain aliphatic aldehydes to fatty acids (Demozay et al., 2008). Such conversion is critical to resolve oxidative stress-induced lipid peroxidation and turnover reactive lipids for their utilization in membrane composition, energy production, or storage. Therefore, we hypothesized that ALH-4 deficient worms might be subject to elevated oxidative stress. As a result, they might be more susceptible to exogenous stress inducers. To this end, we compared wild type and *alh-4(−)* worms for their sensitivity to hydrogen peroxide and paraquat. We found that *alh-4(−)* worms had comparable sensitivity to paraquat as wild type worms (Fig 4A). Surprisingly, *alh-4(−)* worms were more resistant to hydrogen peroxide than wild type worms, and showed enhanced survival after being exposed to 0.75mM hydrogen peroxide for 4 hours (Fig 4B). Our results suggested that *alh-4(−)* worms might mount a homeostatic anti-oxidative response to counter the accumulation of endogenous reactive lipids. As a result, they became more resistant to some but not all exogenous stress inducers as well. To elucidate the molecular basis of such response, we compared the gene expression profiles of wild type and *alh-4(−)* worms by high throughput RNA sequencing.

**Figure 4.**
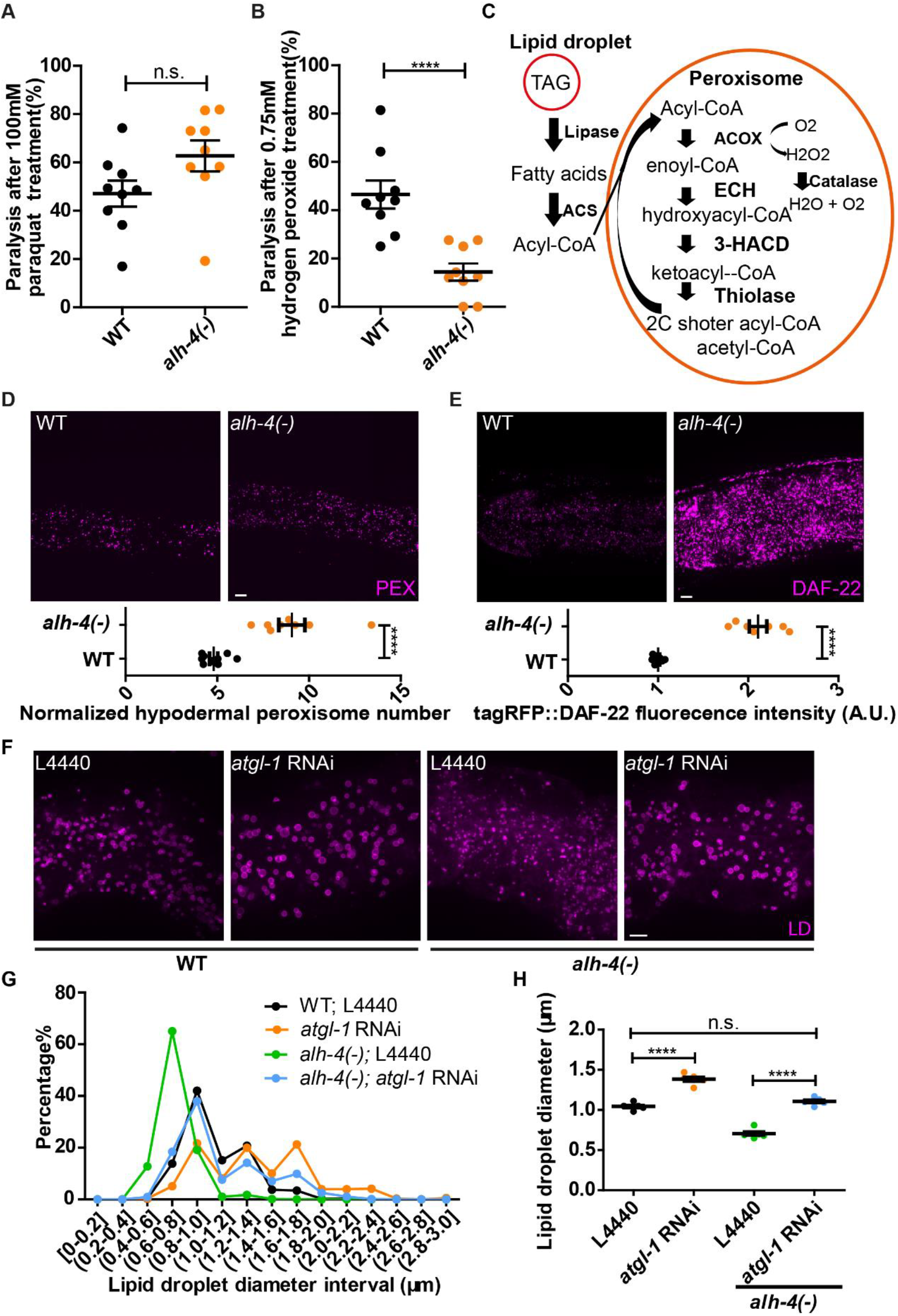
Induction of peroxisomal fatty acid catabolism in *alh-4* mutants. (A) Scatter plot showing the paralysis rate of worms after being treated with 100mM Paraquat for 6 hours. Three independent biological samples of each genotype were tested. Each dot represents one of three technical replicates of each biological sample. (B) As in (A), but showing the paralysis rate of worms after 4 hours treatment with 0.75mM hydrogen peroxide in M9 buffer. (C) Schematic representation of the peroxisomal fatty acid β-oxidation pathway. At the top left corner, triacylglycerols (TAG) in the lipid droplet are hydrolyzed into free fatty acids by lipases. The fatty acids are converted to fatty acyl-CoA by acyl-CoA synthetases. Acyl-CoA oxidases pass electron directly to oxygen and produce hydrogen peroxide (H2O2) while introducing the double bond to fatty acyl-CoA. Hydrogen peroxide is converted to oxygen and water via catalases. ACS, acyl-CoA synthetase; ACOX, acyl-CoA oxidase; ECH, enoyl-CoA hydratase; HACD, hydroxyalkyl-CoA dehydrogenase. (D) Representative images of hypodermal peroxisomes labeled by tagRFP::PTS1 (*hjSi486*) in WT and *alh-4(−)* worms (top). Quantification of hypodermal peroxisome number (bottom). (E) Representative images showing the fluorescence signal of DAF-22 tagged with tagRFP (*hj234*) in WT and *alh-4(−)* worms (top). Quantification of fluorescence signals (bottom). (F) Representative images of lipid droplets labeled with DHS-3::mRuby (*hj200*) in the second intestinal segment. (G) Intestinal lipid droplet size distribution. n = 925 for WT; L4440, n = 526 for *atgl-1* RNAi, n = 672 for *alh-4(−)*; L4440 and n = 706 for *alh-4(−)*; *atgl-1* RNAi. (H) Quantification of lipid droplet diameter in the second intestinal segment. For all statistical analysis, *p < 0.05, **p < 0.01, ***p < 0.001; ****p < 0.0001 (unpaired Student’s t-test). PEX, peroxisome; LD, lipid droplet. Scale bar = 5μm.

We classified statistically significant, differentially expressed genes, using the g:Profiler analysis platform (Reimand et al., 2016). We found that the differentially expressed genes were highly enriched under the KEGG category of ‘peroxisome’ (Fig S4A). This was followed by categories that were broadly related to fatty acid metabolism. Accordingly, a suite of genes that encode enzymes for peroxisomal function, including fatty acid β-oxidation, were significantly upregulated in *alh-4(−)* worms (Fig 4C, Fig S4B). We also noted the concerted induction of *ctl-1*, *ctl-2*, and *ctl-3*, which encode peroxisomal catalases that are known to neutralize hydrogen peroxide, generated during peroxisomal fatty acid β-oxidation (Fig S4B). It is plausible that these catalases may protect *alh-4(−)* worms against exogenous hydrogen peroxide treatment (Fig 4B).

Next, we sought to correlate the upregulation of peroxisome related genes in *alh-4(−)* worms with an increase in peroxisome number or function. We focused on the intestine and hypodermis, where *alh-4* was primarily expressed. To this end, we generated tissue-specific, single-copy transgenic reporter strains, which expressed red fluorescent tagRFP that was targeted to the peroxisomal matrix by the peroxisomal targeting signal PTS1. We found that the density of peroxisomes was significantly increased in the hypodermis, but not the intestine (Fig 4D, S4C). To extend this observation, we focused on *daf-22*, whose expression was induced in *alh-4(−)* worms according to our RNA sequencing data (Fig S4B). We inserted the coding sequence of tagRFP into *daf-22* by CRISPR, to yield transgenic worms that expressed a functional tagRFP::DAF-22 fusion protein from its endogenous locus. We observed an overall 2-fold increase of tagRFP::DAF-22 fluorescence signal in *alh-4(−)* worms (Fig 4E). Taken together, we proposed that the enhanced resistance of *alh-4(−)* worms to hydrogen peroxide was collateral to peroxisome proliferation and the elevation of peroxisomal activity.

### Induction of *atgl-1* in *alh-4* mutant worms

From our gene expression analysis, we noted that *atgl-1*, which encodes the *C. elegans* adipose triglyceride lipase (ATGL) ortholog, was induced in *alh-4(−)* worms. ATGL hydrolyzes a fatty acyl chain from TAG to yield DAG (Young and Zechner, 2013). Fatty acids can then be activated by acyl-CoA synthetases (ACS) to form acyl-CoA before being channeled to anabolic or catabolic pathways, such as β-oxidation. Therefore, the induction of ATGL reduces cellular fat storage. Accordingly, we observed an increase in average intestinal LD size when *atgl-1* was knocked down by RNAi (Fig 4F-H). Furthermore, RNAi against *atgl-1* in *alh-4(−)* worms restored the average LD size to that observed in wild type worms (Fig 4F-H). Our results indicated that the increase in *atgl-1* expression contributed to the reduced fat storage in *alh-4(−)* worms.

### NHR-49 and NHR-79 are required for the transcriptional response to ALH-4 deficiency

To identify transcription factors that are responsible for the altered gene expression program in *alh-4(−)* worms, we focused our bioinformatic analysis on 737 differentially expressed genes. We extracted 1kb sequence 5’ to the start codon of each gene and identified enriched sequence motifs by RSAT. We then queried footprintDB for transcription factors that were predicted to bind these motifs (Sebastian and Contreras-Moreira, 2014), which yielded 42 transcription factors. An additional 10 transcription factors that were reported to regulate fat storage were considered (Arda et al., 2010; Mori et al., 2017; Pathare et al., 2012). Next, we curated a cherry-picked RNAi library of these 52 transcription factors in order to test if their depletion would modulate *alh-4(−)* phenotypes. To this end, we focused on the hypodermal peroxisome density and the fluorescence intensity of tagRFP::DAF-22, expressed from the endogenous *daf-22* locus. Our functional genomic analysis revealed two nuclear receptors, NHR-49 and NHR-79 that are in part responsible for the transcriptional response in *alh-4(−)* worms. To confirm our results based on RNAi, we introduced *nhr-49(lf)* or *nhr-79(lf)* mutant alleles into *alh-4(−)* worms. The loss of NHR-49 or NHR-79 significantly reduced peroxisome density and tagRFP::DAF-22 fluorescence intensity in *alh-4(−)* worms to levels lower than that seen in wild type worms (Fig 5B-E). We detected NHR-49 and NHR-79 expression in the intestine and hypodermis (Fig S5B). It is plausible that NHR-49 and NHR-79 act together in the hypodermis to regulate peroxisome proliferation and function in response to ALH-4 deficiency. Notably, NHR-49 and NHR-79 were reported to interact with each other in yeast two-hybrid assays (Pathare et al., 2012).

**Figure 5.**
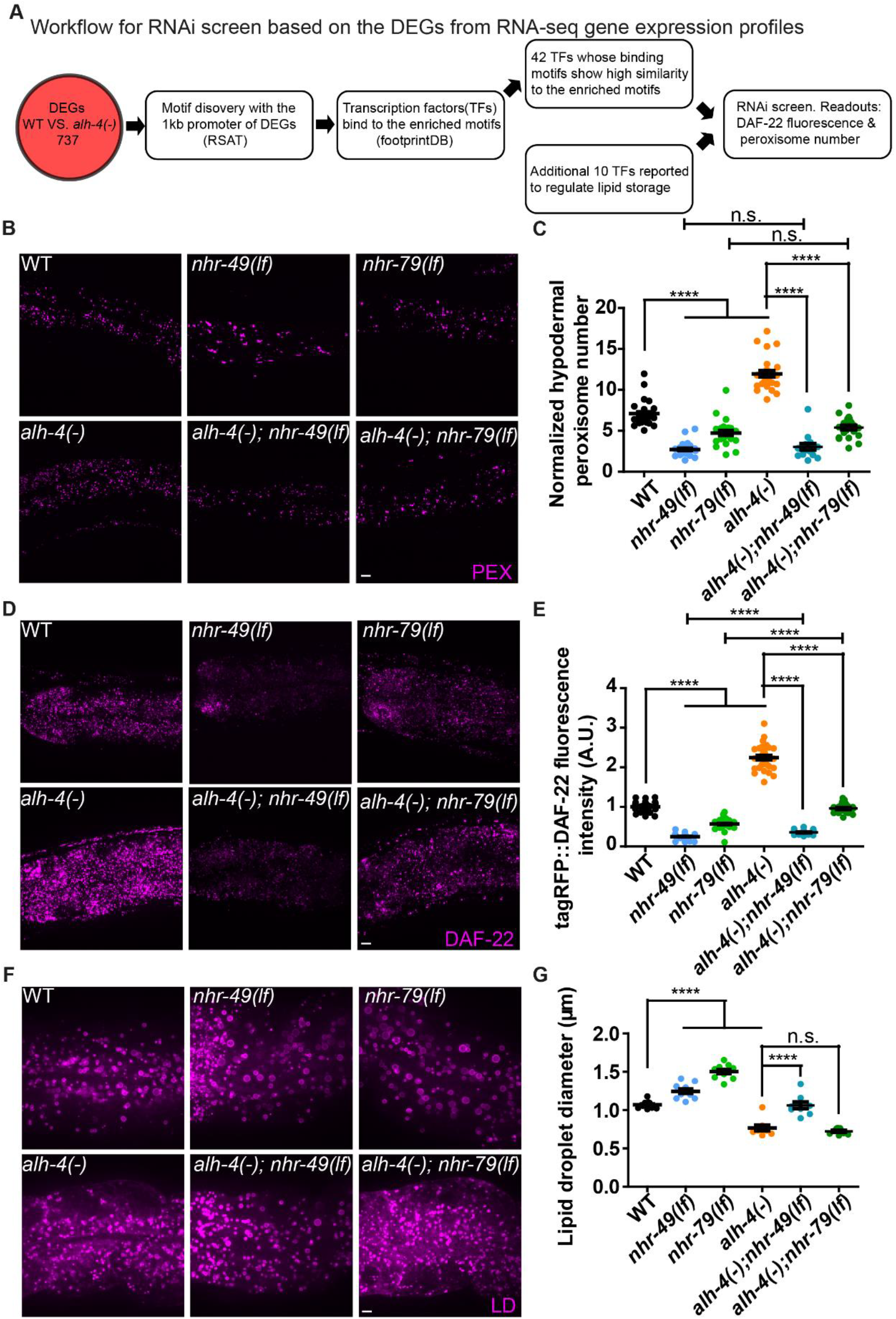
Loss of *nhr-49* or *nhr-79* function suppresses ALH-4 deficiency. (A) Workflow for RNAi screening based on differentially expressed genes (DEGs) identified by RNA sequencing of wild type (WT) and *alh-4(−)* worms. 737 genes were found to be differentially expressed with the threshold fold change >2 and p-value < 0.05. (B) Representative images showing hypodermal peroxisomes labeled by tagRFP::PTS1 *(hjSi486)*, in WT, *nhr-49*(*lf*), *nhr-79*(*lf*), *alh-4(−)*, *alh-4(−)*; *nhr-49(lf)* and *alh-4(−)*; *nhr-79(lf)* L4 worms. (C) Quantification of hypodermal peroxisome number, n> 20 for each genotype from 3 independent experiments. (D) As in (B), but with animals expressing tagRFP::DAF-22 from the endogenous *daf-22* locus. (E) Quantification of tagRFP::DAF-22 fluorescence intensity. n >20 for each genotype from 3 independent experiments. (F) As in (B), but with animals expressing the lipid droplet marker DHS-3::mRuby (*hj200*). (G) Quantification of lipid droplet size in the second intestinal segment. n = 10 for each genotype from 2 independent experiments. In each plot, mean ± SEM of each group is shown. For all statistical analysis, *p < 0.05, **p < 0.01, ***p < 0.001; ****p < 0.0001 (unpaired Student’s t-test). PEX, peroxisome; LD, lipid droplet. Scale bar = 5μm.

Next, we asked if NHR-49 and NHR-79 were also responsible for mounting a transcriptional response to ALH-4 deficiency in the intestine. The intestinal LDs in *nhr-49(lf)* or *nhr-79(lf)* mutant worms was significantly larger than those in wild type worms (Fig 5F-G). However, the loss of NHR-49 but not NHR-79 corrected the intestinal LD size of *alh-4(−)* worms (Fig 5F-G). Therefore, NHR-49 and NHR-79 appeared to act independently in the intestine to control LD size: only NHR-49 was responsible for the transcriptional response to ALH-4 deficiency.

### Intestinal ALH-4 regulates hypodermal peroxisome proliferation

The differential suppression of ALH-4 deficiency in *nhr-49(lf)* and *nhr-79(lf)* mutant worms suggested that tissue specific transcriptional programs may operate to sense cellular fatty aldehyde levels. If so, is organismal fatty aldehyde metabolism coordinated between tissues? To this end, we first sought to determine the site of action of ALH-4. We generated two single-copy transgenes that specifically expressed functional GFP::ALH-4 fusion proteins in the intestine or hypodermis of *alh-4(−)* worms (Fig S6A). The restoration of *alh-4* expression, as indicated by GFP fluorescence in the intestine, normalized the intestinal LD size and tagRFP::DAF-22 expression to wild type level (Fig 6A and 6C). Furthermore, the hypodermal peroxisome density and tagRFP::DAF-22 expression of these worms were corrected to the wild type level (Fig 6B). In contrast, hypodermis-specific restoration of ALH-4 expression in *alh-4(−)* worms failed to normalize intestinal LD size, hypodermal peroxisome density, and tagRFP::DAF-22 expression level in both tissues (Fig 6A-C). Our results indicate that intestinal ALH-4 regulates intestinal lipid storage cell-autonomously and hypodermal peroxisome proliferation cell-non-autonomously. In *alh-4(−)* worms, it is plausible that the alteration of cellular fatty aldehyde levels yields signals that act on the intestine and hypodermis in an autocrine and endocrine fashion, respectively.

**Figure 6.**
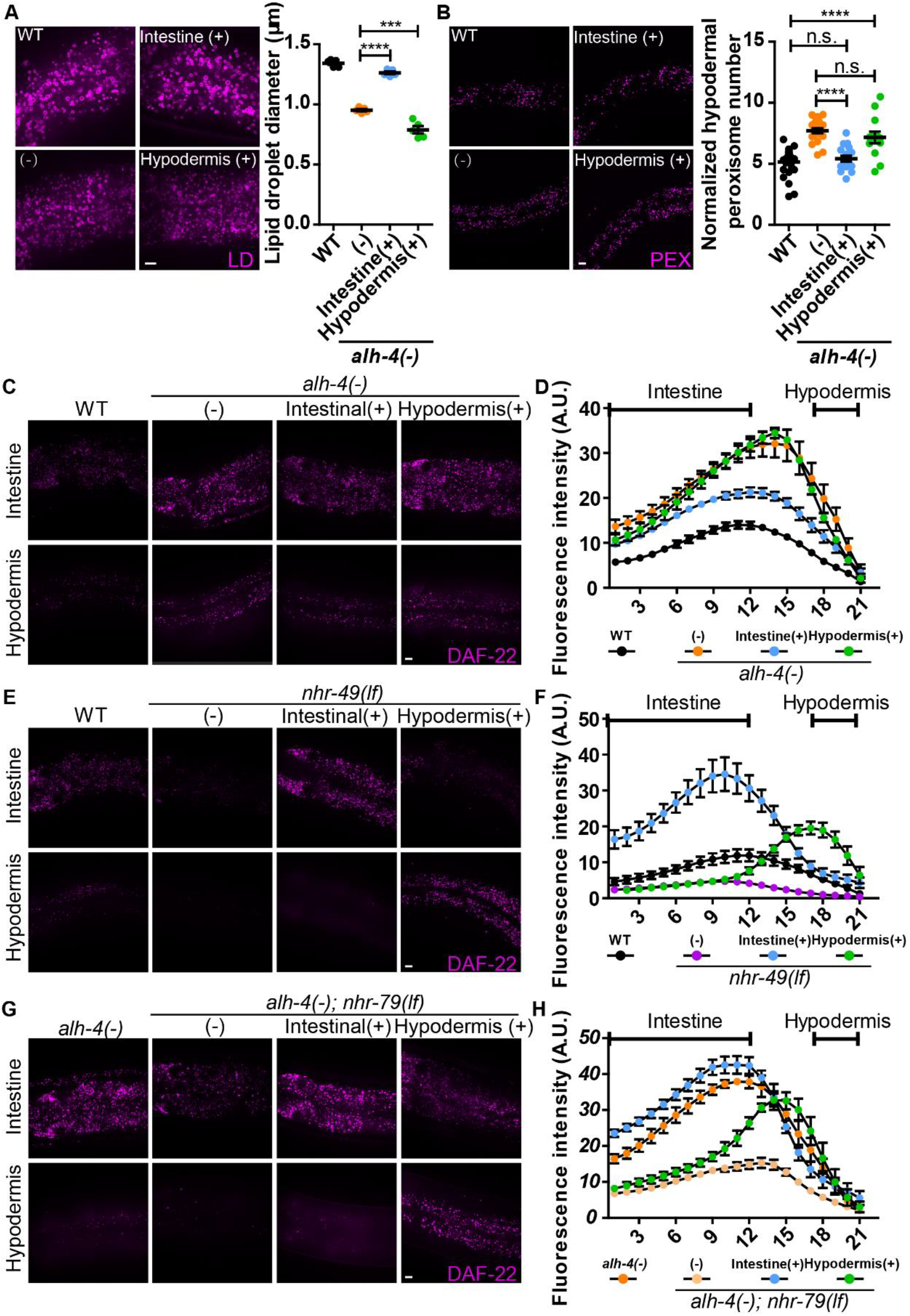
Tissue specific rescue of *alh-4*, *nhr-49* and *nhr-79* mutant animals. (A) Representative images showing intestinal lipid droplets labeled with DHS-3::mRuby (*hj200)* (left). Quantification of lipid droplet diameter in the second intestinal segment (right). n = 5 for each group. (B) Representative images showing hypodermal peroxisomes labeled with tagRFP::PTS1 (*hjSi486)* (left). Quantification of hypodermal peroxisomes normalized with the length of the worm in the field of view (right). n > 13 for each group. (C) Representative images of Z-stack of 3 focal planes at 0.5μm interval showing tagRFP::DAF-22 in the intestine (top) and hypodermis (bottom) in wild type (WT), *alh-4(−)*, *alh-4(−); hjSi501[vha-6p::alh-4(+)]*, *alh-4(−); hjSi500[dpy-7p::alh-4(+)]* worms. (D) Quantification of fluorescence intensity of tagRFP::DAF-22 in each focal plane of Z stacks that encompass the intestine and hypodermis. n = 6 for each group. (E) As in (C), but with wild type (WT), *nhr-49(lf)*, *nhr-49(lf); hjSi531[vha-6p:: nhr-49(+)]*, *nhr-49(lf); hjSi537[dpy-7p:: nhr-49(+)]* worms. (F) As in (D), but with worms of specified genotypes. n = 5 for WT and n = 6 for other groups (G) As in (C), but with wild type (WT), *alh-4(−); nhr-79(lf)*, *alh-4(-); nhr-79(lf); hjSi553[vha-6p:: nhr-79(+)]*, *alh-4(−); nhr-79(lf); hjSi539[dpy-7p:: nhr-79(+)]* worms. (H) As in (D), but with worms of specified genotypes. n = 6 for each group. For all statistical analysis, *p < 0.05, **p < 0.01, ***p < 0.001; ****p < 0.0001 (unpaired Student’s t-test). PEX, peroxisome; LD, lipid droplet. Scale bar = 5μm.

### NHR-49 and NHR-79 act cell-autonomously in response to ALH-4 deficiency

Next, we determined the site of NHR-49 and NHR-79 action. We constructed transgenic worms that expressed GFP fusion of NHR-49 or NHR-79 specifically in the intestine or hypodermis from single copy transgenes, under the control of *vha-6* or *dpy-7* promoter, respectively. The tissue specific expression of fusion proteins was confirmed by the detection of primarily nuclear fluorescence signals in the appropriate tissues (Fig S6B-C). We found that NHR-49 acted cell-autonomously in the intestine and hypodermis. Accordingly, intestinal expression of NHR-49 normalized tagRFP::DAF-22 expression and LD size in the intestine but not hypodermis (Fig 6E-F, S6E). Hypodermal expression of NHR-49 specifically normalized hypodermal peroxisome density and tagRFP::DAF-22 expression and had no impact on mutant phenotypes in the intestine (Fig 6E-F, S6D). Similar observations were made for NHR-79: restoration of intestinal and hypodermal NHR-79 expression specifically suppressed mutant phenotypes in the respective tissues in a cell-autonomous manner (Fig 6G and S6F). Taken together, we conclude that NHR-49 and NHR-79 act cell-autonomously in the intestine and hypodermis.

### NHR-49 functionally couples with NHR-79 to regulate peroxisome proliferation

Most nuclear receptors function as homodimers or heterodimers. NHR-49 partners with NHR-66, NHR-80 and NHR-13 to regulate distinct aspects of lipid metabolism (Pathare et al., 2012). NHR-49 was also reported to interact with NHR-79 in yeast two-hybrid interaction assays (Pathare et al., 2012). Therefore, we asked if NHR-49 and NHR-79 were functionally coupled to regulate fat storage and peroxisome proliferation in response to ALH-4 deficiency. First, we focused on the expression of the tagRFP::DAF-22 fusion protein that was expressed from the endogenous *daf-22* locus. In mutant worms that lacked NHR-49 or NHR-79, the expression of tagRFP::DAF-22 expression was reduced (Fig 7A-B). No further reduction was observed when both *nhr-49* and *nhr-79* were mutated (Fig 7A-B). Therefore, NHR-49 and NHR-79 appeared to act in concert to regulate *daf-22* expression. Our conclusion was further substantiated by the following observation: the elevation of *daf-22* expression in *nhr-49* gain-of-function (*gf*) mutant worms was partially dependent on *nhr-79* (Fig 7A-B). NHR-49 and NHR-79 appeared to act in a similar manner to regulate hypodermal peroxisome density (Fig 7A and C). Next, we measured the intestinal lipid droplet size in *nhr-49* and *nhr-79* mutant worms. NHR-79 deficiency caused a larger increase in lipid droplet size than NHR-49 deficiency (Fig 7A and D). Interestingly, the lipid droplet size in *nhr-49(lf)* and *nhr-79(lf)*; *nhr-49(lf)* mutant worms showed no significant difference. In addition, gain of NHR-49 function reduced the lipid droplet size of *nhr-79(lf)* worms (Fig 7A and D). Our results suggest that NHR-49 and NHR-79 regulate distinct sets of genes in the intestine, which modulate lipid droplet size independently. Taken together, NHR-49 and NHR-79 act together to regulate genes pertaining to peroxisome number and function. However, each of them may have additional functional partners to control fat storage in the intestine.

**Figure 7.**
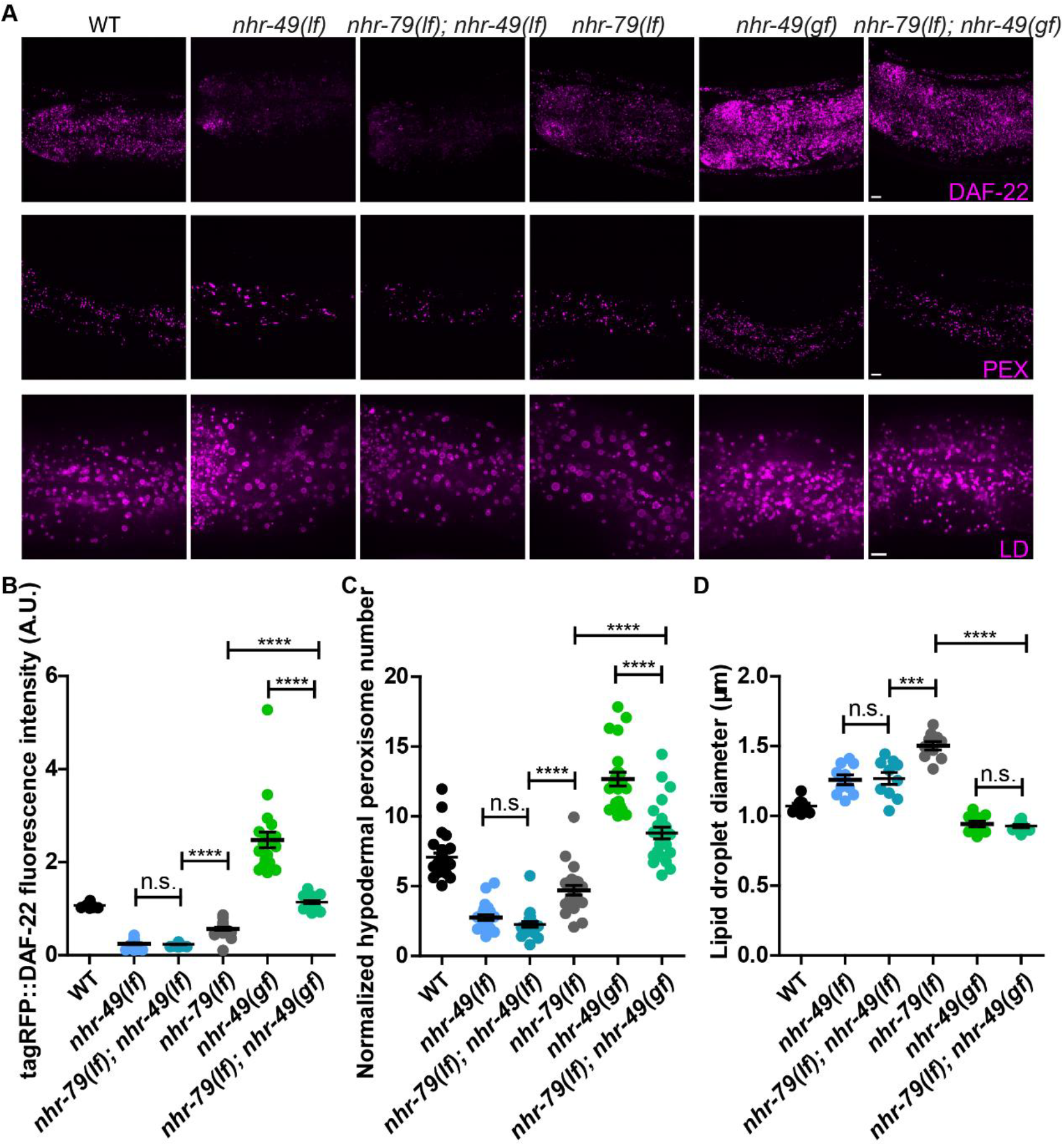
Functional interaction of NHR-49 and NHR-79. (A) Representative images showing the fluorescence of tagRFP::DAF-22 (*hj234*) (top), hypodermal peroxisomes labeled with tagRFP::PTS1(*hjSi486*) (middle), and intestinal lipid droplets labeled with DHS-3::mRuby (*hj200*) (bottom). (B) Quantification of fluorescence intensity of tagRFP::DAF-22. n > 20 for each group. (C) Quantification of hypodermal peroxisomes normalized with the length of the worm in the field of view. n > 20 for each group. (D) Quantification of lipid droplet size in the second intestinal segment. n = 10 for each group. In each plot, mean ± SEM of each group is shown. For all statistical analysis, *p < 0.05, **p < 0.01, ***p < 0.001; ****p < 0.0001 (unpaired Student’s t-test). PEX, peroxisome; LD, lipid droplet. Scale bar = 5μm.

### NHR-49 is required for the reproduction of ALH-4 deficient worms

Despite the accumulation of fatty aldehydes (Fig 1F) and presumably their toxic adducts, *alh-4(−)* worms did not exhibit heightened sensitivity to oxidative stress (Fig 4A-B). We reasoned that the enhanced resistance of *alh-4(−)* worms to hydrogen peroxide was due to NHR-49/NHR-79 dependent peroxisome proliferation and the concurrent overexpression of catalases (Fig 4C). Therefore, peroxisome proliferation could be regarded as a compensatory response that maintained the health of *alh-4(−)* worms. To test this hypothesis, we used the number of progenies as an indicator of normal reproductive development. Both *alh-4(−)* and *nhr-49(lf)* mutant worms had a ~30% reduction of brood size (Fig 8A). However, almost all *alh-4(−)*; *nhr-49(lf)* double mutant worms failed to produce live progeny after they developed into adults (Fig 8A). These genetically lethal worms could be rescued by an extrachromosomal array that carried the wild type *alh-4* gene (Fig 8A). In contrast, the reproduction of *alh-4(−)* worms was not strictly dependent on NHR-79. Accordingly, *alh-4(−)*; *nhr-79(lf)* mutant worms remained fertile, although their brood size was reduced in comparison to that of *alh-4(−)* or *nhr-79(lf)* single mutant worms (Fig 8A). We conclude that NHR-49 is a key factor that supports reproductive development of *alh-4* deficient worms, in part because of its broad role in regulating peroxisome proliferation and fat storage.

**Figure 8.**
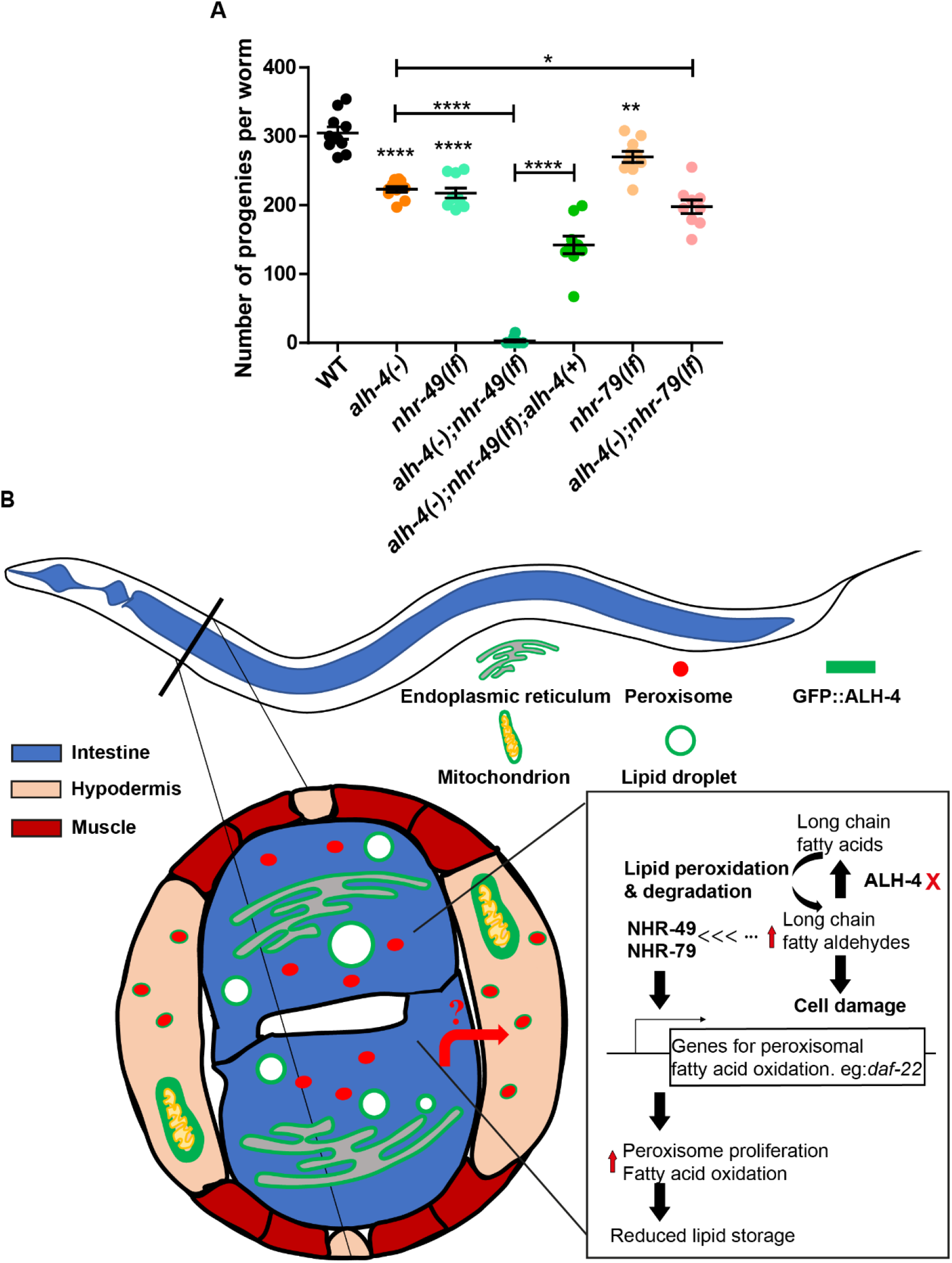
NHR-49 is required for the fertility of *alh-4(−)* worms. (A) Quantification of brood size of wild type (WT), mutant worms of indicated genotypes. n > 8 for each group. In the plot, mean ± SEM of each group is shown. For all statistical analysis, *p < 0.05, **p < 0.01, ***p < 0.001; ****p < 0.0001 (unpaired Student’s t-test). (B) A model for NHR-49/NHR-79 dependent peroxisome proliferation in response to ALH-4 deficiency.

## Discussion

In this paper, we combined genetic, imaging, and gene expression profiling approaches to identify a peroxisome proliferative response that compensated for ALH-4 deficiency in *C. elegans* (Fig 8B). We observed concerted induction of genes that encode enzymes for TAG lipolysis and peroxisomal fatty acid catabolism, which contributed to the reduction of neutral lipid storage in *alh-4* mutant worms. In addition, upregulation of peroxisomal catalases in these worms conferred heightened resistance to exogenous hydrogen peroxide, an inducer of oxidative stress. Functional genomic analysis identified NHR-49 and NHR-79 as key transcription factors that were required for peroxisome proliferation in *alh-4* mutant worms. Interestingly, NHR-49 and NHR-79 acted cell autonomously in the hypodermis in response to ALH-4 deficiency in the intestine, thus hinting at an unknown endocrine signal that facilitates intestine to hypodermis communication. Our work revealed an unexpected homeostatic mechanism that alleviates fatty aldehyde dehydrogenase deficiency.

The loss of FALDH function causes Sjögren-Larsson Syndrome in humans (De Laurenzi et al., 1996). The prominent phenotypes of spasticity and ichthyosis led to the proposal that the nervous system and skin are particularly reliant on FALDH for the metabolism of sphingolipids and the resolution of fatty aldehydes. The pathology of Sjögren-Larsson Syndrome has been partially recapitulated in FALDH/*Aldh3a2* knockout mice (Kanetake et al., 2019; Naganuma et al., 2016). Detailed biochemical analysis of lipid species from skin and brain tissues of these knockout mice identified specific alteration in sphingolipid intermediates, which may contribute to keratinocytes and neuronal dysfunction. In this paper, we present an invertebrate model of FALDH deficiency in the form of *alh-4* mutant worms. Based on sequence homology and the characteristic membrane association,

ALH-4 is likely the sole ortholog of FALDH in *C. elegans*. As such, ALH-4 deficient worms have excess fatty aldehydes or fatty alcohols (Fig 1F, S1G), similar to FALDH deficient fibroblasts (Rizzo and Craft, 2000). However, we did not detect gross neuronal and epidermal defects based on available assays. This may be attributed to fundamental differences in tissue architecture. For example, it has been hypothesized that myelin dysfunction contributes to neural symptoms in Sjögren-Larsson Syndrome (Kanetake et al., 2019). However, *C. elegans* neurons are not myelinated.

We originally cloned *alh-4* based on the observation that *alh-4* loss of function alleles suppressed fat accumulation and LD expansion in *daf-22* mutant worms. Interestingly, the expression of DAF-22, together with other peroxisomal β-oxidation enzymes, was induced in *alh-4* mutant worms. In contrast, the expression of genes encoding mitochondrial β-oxidation enzymes including acyl-CoA synthetases (ACSs), Enoyl-CoA Hydratases (ECHs), Carnitine Palmitoyl Transferases (CPTs), Acyl CoA Dehydrogenases (ACDHs) and thiolases were relatively unperturbed. Our results are consistent with the notion that *alh-4* deficiency specifically activates peroxisomal fatty acid β-oxidation pathways that act in parallel of the DAF-22 pathway. Therefore, enhanced fatty acid catabolism in the absence of ALH-4 prevented neutral fat storage in *daf-22* mutant worms.

By engineering worms that expressed a cytosolic form of ALH-4, we confirmed that membrane association is essential for its proper function. ALH-4 and mammalian FALDH share the same mechanism of using divergent C-termini for tail-anchored organelle-specific association (Ashibe et al., 2007; Costello et al., 2017). Therefore, it is conceivable that ALH-4 processes fatty aldehydes in situ as they arise in organelle membranes, thus necessitating the distribution of ALH-4 isoforms to multiple organelles. Surprisingly, we found that each of the ALH-4 isoforms was sufficient to rescue *alh-4(−)* mutant worms. For example, the expression of ALH-4A, which associates with mitochondria and peroxisomes, relieved the requirement of ALH-4B and ALH-4C on the ER and LDs. It is formally possible that alternative aldehyde dehydrogenases substituted for ALH-4 at the ER and LDs in this case. Nevertheless, our results raised the possibility that fatty aldehydes or their precursors, as part of a membrane lipid, could be transferred from one organelle to another, perhaps via inter-organelle contact sites.

The nuclear receptor NHR-49 partners with NHR-66, NHR-80 and NHR-13 to regulate distinct sets of target genes in lipid catabolism, desaturation and remodeling pathways (Pathare et al., 2012). Based on quantitative proteomic analysis, there is a good correlation between differentially expressed genes and their corresponding proteins in *nhr-49* knock-down worms (Fredens and Færgeman, 2012). Although NHR-49 has long been proposed as a functional ortholog of PPARα (Van Gilst et al., 2005b, 2005a), the direct demonstration of its ability to promote peroxisome proliferation has been elusive. Here, we show that NHR-49, in partnership with NHR-79, is necessary for peroxisome proliferation in wild type and *alh-4* mutant worms. Interestingly, although NHR-49 and NHR-79 are both expressed in the intestine and hypodermis, the mRNA level of NHR-49 is >150 times higher than NHR-79 in wild type and *alh-4* mutant worms (Fig S5A). In comparison, the expression level of NHR-49 is 5 to 48 times higher than NHR-66, NHR-80 or NHR-13. This is consistent with the notion that NHR-49 serves as a common partner to multiple nuclear receptors in a tissue or context dependent manner. Accordingly, NHR-49 acts with NHR-79 to promote peroxisome proliferation in *alh-4* mutant worms. However, NHR-49 and NHR-79 regulate LD size in a divergent manner (Fig 5F-G).

It has been reported that phospholipids and their metabolites can serve as ligands for *C. elegans* NHRs (Folick et al., 2015; Lin and Wang, 2017; Mullaney et al., 2010). For example, oleoylethanolamide, a PPARα agonist, binds NHR-80 to regulate *C. elegans* aging (Folick et al., 2015; Fu et al., 2003). Although the polyunsaturated α-linolenic acid was reported to act via NHR-49 to extend *C. elegans* lifespan (Qi et al., 2017), no cognate ligand has been identified for NHR-49 or NHR-79 through direct binding assays. Interestingly, exogenous organic peroxide was reported to activate an NHR-49 dependent stress resistance program (Goh et al., 2018). Taken together, it is tempting to speculate that excess fatty aldehydes or their metabolites in *alh-4* mutant worms may activate NHR-49 or NHR-79. Furthermore, the putative activators should be able to travel from the intestine to the hypodermis. This is because intestinal ALH-4 deficiency can trigger NHR-49/NHR-79 dependent peroxisome proliferation in the hypodermis. To substantiate our hypothesis, further analysis of metabolites that are enriched in *alh-4* mutant worms will be necessary.

The concept of neurons-glia communication via lipid metabolites has been advanced in specific forms of neurodegeneration (Chung et al., 2020; Liu et al., 2017). One therapeutic approach focused on the reduction of ROS, which are regarded as the upstream trigger of cell damage (Chung et al., 2020). In principle, the modulation of latent, endogenous pathways may offer alternative therapeutic strategies. It is unclear if cell-cell communication also underlies the pathology of Sjögren-Larsson Syndrome. Our work indicates that an NHR-49 dependent, peroxisome proliferation and lipid detoxification program is required for the health and propagation of ALH-4 deficient worms. By analogy, fine-tuning of peroxisome proliferation or function via PPAR activation may be a probable strategy for treating Sjögren-Larsson Syndrome. Notably, PPARα induces FALDH expression (Ashibe and Motojima, 2009) and the PPARα agonist Bezafibrate has been suggested as an agent for treating Sjögren-Larsson Syndrome patients with partially inactivated FALDH (Gloerich et al., 2006). Additional natural and synthetic PPAR ligands have been actively tested as potential treatments for neurologic disorders (Fidaleo et al., 2014; Moutinho et al., 2019). It is conceivable that the same ligands can be repurposed to benefit Sjögren-Larsson Syndrome patients.

## Materials and Methods

### *C. elegans* strains and maintenance

*C. elegans* strains are fed with *E. coli* OP50 on standard Nematode Growth Medium (NGM) plates at 20℃ using standard protocol. Bristol N2 was used as the wild type. Strains carrying the following alleles were originally obtained from the *Caenorhabditis* Genetics Center (CGC): LG II, *daf-22*(*ok693*); LG V, *nhr-79*(*gk20*); LG I, *nhr-49*(*nr2041*), *nhr-49*(*et8*).

Mutant alleles first reported in this study are as follows. LGV, *alh-4(hj27)*, *alh-4(hj28)*, *alh-4(hj29)*, *alh-4(hj221)*

Knock-in strains obtained with CRISPR-SEC (Dickinson et al., 2015):

*alh-4(hj179)* [*GFP_TEV_loxP_3XFLAG_linker_alh-4*],

*dhs-3(hj200)* [*dhs-3_linker_mRuby_TEV_3XFLAG*],

*daf-22(hj234)* [*tagRFP_TEV_loxP_3xFLAG_linker_daf-22*],

*alh-4(hj261)* [*GFP_TEV_loxP_3XFLAG_linker_alh-4(1-463aa)*],

*alh-4 (hj264)* [*GFP_TEV_loxP_3XFLAG_linker_alh-4(449-459aa deletion)*],

*nhr-49(hj293)* [*nhr-49_linker_GFP_TEV_3xHA*].

Single-copy transgenes obtained with MosSCI (Frøkjær-Jensen et al., 2012) or CRISPR-SEC with a sgRNA targeting Mos1 (from Jihong Bai): *hjSi112*[*vha-6p::3xFLAG::TEV::mRuby::F59A1.10 coding::let-858 3’UTR*] *IV, hjSi348*[*vha-6p::acs-22 cDNA::tagRFP::TEV::3xFLAG::let-858 3’ UTR*] *IV, hjSi486*[*dpy-7p::tagRFP::PTS1::let-858 3’ UTR*] *IV, hjSi489*[*dpy-30p::tomm-20(1-54aa)::mRuby::let-858 3’ UTR*] *II, hjSi495*[*dpy-30p::GFP::alh-4c(454-494aa)::let-858 3’ UTR*] *I, hjSi497*[*dpy-30p::GFP::alh-4a(454-493aa)::let-858 3’ UTR*] *I, hjSi499*[*dpy-30p::GFP::alh-4b(454-493aa)::let-858 3’ UTR*] *I, hjSi500*[*dpy-7p::GFP::TEV::loxP::3xFLAG::alh-4::dhs-28 3’UTR*] *I, hjSi501*[*vha-6p::GFP::TEV::loxP::3xFLAG::alh-4::dhs-28 3’UTR*] *I, hjSi506*[*dpy-7p::GFP::PTS1:: let-858 3’ UTR*] *IV, hjSi531*[*vha-6p::nhr-49(a,b isoform cDNA with one short intron)::GFP::HA::dhs-28 3’UTR*] *II, hjSi537*[*dpy-7p::nhr-49(a, b isoform cDNA with one short intron)::GFP::HA::dhs-28 3’UTR*] *II, hjSi539*[*dpy-7p::GFP::3xFLAG::nhr-79a isoform(cDNA)::dhs-28 3’UTR*] *IV, hjSi548*[*vha-6p::mRuby::PTS1::dhs-28 3’ UTR*] *IV, hjSi553*[*vha-6p::GFP::3xFLAG::nhr-79a isoform(cDNA)::dhs-28 3’UTR*] *II, hjSi554*[*dpy-30p::GFP::TEV::3xFLAG::alh-4a::let-858 3’ UTR*] *I, hjSi555*[*dpy-30p::GFP::TEV::3xFLAG::alh-4b::let-858 3’ UTR*] *I, hjSi556*[*dpy-30p::GFP::TEV::3xFLAG::alh-4c::let-858 3’ UTR*] *I.*

Extra-chromosomal array strains:

*hjEx25[dpy-30p::GFP::TEV::3xFLAG::alh-4::let-858 3’ UTR in pCFJ352]*

*hjEx26[nhr-79p::mRuby::3xFLAG::nhr-79::nhr-79 3’UTR; tub-1p::GFP::tub-1]*

All strains had been outcrossed with wild type N2 at least twice before being used.

### *C. elegans* genetic screen

The *daf-22(ok693)* mutant worms were mutagenized with Ethyl Methane Sulfonate (EMS) for suppressors of the expanded LD phenotype. Three mutant alleles (*hj27*, *hj28* and *hj29*) were mapped to *alh-4* using a single nucleotide polymorphism–based strategy with the Hawaiian C. elegans isolate CB4856 (Davis et al., 2005). Molecular lesions of mutant alleles were determined by targeted Sanger sequencing. *hj27* encodes a G to A mutation that causes the substitution of Gly 415 to Glu. *hj28* and *hj29* are nonsense alleles. *hj28* carries a C to T substitution (Gln 301 stop), and *hj29* carries a G to A substitution (Trp 236 stop).

### RNA interference-based knockdown in *C. elegans*

The RNA interference (RNAi) constructs were transformed into a modified OP50 strain that is competent for RNAi (Neve et al., 2020). Details of RNAi clones for transcription factors are available in the supplementary data 1 sheet 5. The target sequence in RNAi clones for *alh-4* and *atgl-1* were amplified by the following primer pairs: [5’-AGTTGTACTCATCATCTCCCCATG-3’, 5’-TCCTCGCCATAAAATTCGTTC-3’] and [5’-TGGAGCATCTTCGTCTTCCTC-3’, 5’-AAAATCATATAAATTTTCAGC-3’], respectively from genomic DNA. OP50 RNAi clones were streaked on LB plates with 100μg/mL ampicillin and tetracycline and cultured with 2xYT with 100μg/mL ampicillin overnight. Fresh liquid culture was seeded onto NGM plates for RNAi. The seeded plates were incubated at room temperature for 1 day and 10 L4 worms were transferred on the seeded RNAi plates. When worms reached day 1 adult, they were transferred to new RNAi plates to lay eggs for overnight and then removed from the plates. F1 worms at L4 stage were used for imaging.

### Hydrogen peroxide treatment

Synchronization was done by collecting L1 worms that were hatched within a 2-hour window. 200 L1 worms were transferred to 60mm NGM plates seeded with OP50. After 48 hours, L4 worms were collected with M9, transferred to 1.5mL Eppendorf tubes and washed three times with M9 buffer and then aliquoted into 2 tubes, one for experimental group and the other for negative control. For experimental groups, M9 buffer with hydrogen peroxide was added to worms (final concentration = 0.75mM). For negative control, M9 without hydrogen peroxide was added. Worms were aliquoted into 24 well plates, about 20 worms per well and 3 wells for worms collected from 1 plate and then incubated at 20°C for 4 hours. To stop the hydrogen peroxide treatment, OP50 bacteria liquid culture was added. Worms were scored after a recovery period of 30 minutes. Worms that did not show detectable movement were categorized as paralyzed. Paralysis rate = (number of the paralyzed worms / total number of worms) _experimental group_ − (number of the paralyzed worms / total number of worms) _negative control group_.

### Paraquat treatment

The procedure for paraquat treatment is the same as that for hydrogen peroxide treatment except that the worms were incubated in M9 with 100mM paraquat instead of hydrogen peroxide.

### Brood size

Individual L4 worms were introduced onto NGM plates seeded with OP50 and transferred to new plates every day until they stopped egg laying. 1.5 days after the mother worms were removed, the number of larvae on each plate was counted. The total number of progeny from each adult worm was calculated and recorded as the brood size.

### Growth rate

50-100 adult worms were transferred to NGM plate seeded with OP50 to lay eggs for 1 hour and then 50 eggs were transferred to each new plate. 70 hours after egg laying, larvae were classified as early L4, mid L4, the stage between L4 and adult or adult stage based on the valval morphology.

### Pharyngeal pumping rate

L4 worms were transferred onto new NGM plates seeded with OP50, 2 worms per plate. After 24 hours, the pharyngeal pumping of 1-day old adults were video recorded on a dissecting microscope for at least 1min at room temperature. The videos were played with the PotPlayer at 0.3X speed for precise counting of pharyngeal muscle contractions per min.

### Fluorescence imaging of *C. elegans*

Fluorescence images were obtained on a spinning disk confocal microscope (AxioObeserver Z1, Carl Zeiss) equipped with a Neo sCMOS camera (Andor) controlled by the iQ3 software (Andor). The excitation lasers for GFP, tagRFP/ mRuby were 488nm and 561nm and the emission filters were 500-550nm and 580.5-653.5nm, respectively. The images were imported to Imaris 8 (Bitplane) for image processing and analysis and exported as files in ims format. The ims files were further imported into ImageJ for analysis.

### The quantification of lipid droplet size

Lipid droplets in the intestinal cells were labeled with fluorescent protein tagged resident lipid droplet proteins. Fluorescence images of L4 larvae were acquired on a spinning disk confocal microscope with a 100x objective. For each worm, a Z-stack was taken at 0.5μm interval and 10μm thickness (21 layers) starting at the layer between the intestinal cell and the hypodermis. From each stack, 10 continuous focal planes were chosen to construct a 3D image. The Z-stack of bright field images was used to determine the outline of the intestinal cells. The lipid droplets in the second intestinal segment were labeled manually with the spot function in the Imaris 8 (Bitplane) and the diameter of each lipid droplet were exported and used to calculate the average diameter of lipid droplets for each worm. When calculating the average lipid droplet diameter, lipid droplets with diameter < 0.5 micrometer were excluded because the diffraction limit of light microscopy preluded their accurate measurement.

### The quantification of peroxisome number

Peroxisomes were labeled with fluorescent protein tagged with a PTS1 signal peptide. Fluorescence images of L4 larvae were acquired on a spinning disk confocal microscope with a 63x objective. For each worm, a Z-stack of its anterior region without the head was taken at 0.5μm interval and 4.5μm thickness, focusing on the tissue of interest. A Z-stack of bright field images was also obtained. Z-stack images were imported to Imaris 8 (Bitplane) for 3D reconstruction. The peroxisomes were automatically detected by the spot function with the following algorithm: Enable Region of interest = false; Enable region growing = true; Source channel index = 2; Estimated Diameter = 0.5; Background Subtraction = true; ‘quality’ above 1.000; Region Growing type = Local contrast; Region Growing automatic Threshold = false; Region Growing manual Threshold = 9; Region Growing Diameter = diameter from volume; Create region channel = false. Spots representing mislabeled peroxisomes or the peroxisomes not in the tissue of interest were manually deleted. For the quantification of the hypodermal peroxisomes, only the peroxisomes in the hypodermis close to the coverslip were counted. With the statistics function, the number of peroxisomes can be obtained. The length of the worm body in the view was measured based the bright field images. The peroxisome number was normalized by the length of the worm body in the field of view.

### The quantification of DAF-22 fluorescence intensity

To measure the expression level of DAF-22, the coding sequencing of tagRFP was inserted at the 5’ end of the endogenous *daf-22* gene by CRISPR, to yield transgenic worms that expressed tagRFP::DAF-22 fusion protein at the endogenous level. Fluorescence images of L4 larvae were acquired on a spinning disk confocal microscope with a 63x objective. For each worm, a Z-stack was taken with 0.5μm interval and 10μm thickness starting at the layer closest to the coverslip with obvious fluorescence signals (i.e. hypodermis) and ending at intestinal layers. A Z-stack of bright field images was also obtained. The field of view included the anterior region of each worm without the head. Using Imaris 8 (Bitplane), the outline of the worm body was manually drawn based on bright field images, the surface function was used to create a worm surface which colocalizes with the surface of the worm. Similarly, a background surface was manually drawn and created in the region of the field without the worm. The average intensity of tagRFP::DAF-22 is the average intensity inside of the worm body surface minus that of the background surface. The normalization of fluorescence intensity was done with the average intensity of WT worms that were imaged at the same time.

### RNA-seq of *C. elegans*

Three biological replicates of N2 and *alh-4(−)* were synchronized and grown to L4 stage. Sequencing library preparation was done according to a published method (Serra et al., 2018) with the modifications: the number of worms for each replicate and the reagent for worm lysis were scaled up by 10 fold and the worms were broken with a mini-pestle in a 1.5ml disposable tube on ice instead of cutting. The libraries were pooled and sequenced with the Illumina NextSeq 500 system. The reads were mapped to ce10 genome assembly. The ce10 Illumina iGenome gene annotation was used as a reference for transcripts identification. The downstream gene expression level analysis was done based on the published method (Pertea et al., 2016) to get FPKM (Fragments Per Kilobase Million) values for differential gene expression analysis.

### Motif analysis

The list of the differentially expressed genes between WT and *alh-4(−)* is obtained with the threshold fold change >2 and p-value < 0.05 (negative binomial test). −1000 to −1 sequence relative to the start codon of differentially expressed genes (DEGs) were obtained for motif discovery to identify enriched motifs with RSAtool (Medina-Rivera et al., 2015). FootprintDB was used to identify transcription factors that are predicted to bind the enriched motifs (Sebastian and Contreras-Moreira, 2014).

### Gene Ontology and pathway analysis

The analysis was done with gProfiler (Reimand et al., 2016) using the list of DEGs between WT and *alh-4(−)* as the input and choosing *Caenorhabditis elegans* as the organism.

### SRS for lipid content measurement

To measure intestinal lipid content, CH_2_ signal was imaged at 2863.5 cm^−1^. Live worms at L4 stage were mounted on 8% agarose pad in PBS buffer with 0.2mM levamisole. The focal plane with maximal SRS signals was determined with a 20x air objective (Plan-Apochromat, 0.8 NA, Zeiss). The quantification of the SRS signal was done following a published protocol (Ramachandran et al., 2015). In short, the average SRS signal intensity in the intestine region per pixel was calculated with the background signal subtracted. The signal intensity for each worm was normalized with the average value of the WT group for the data display.

### Lipid extraction

Lipid were extracted according to a modified version of the methyl-tertiary-butyl ether (MTBE) extraction originally developed by Matyash *et al* (Matyash et al., 2008; Witting et al., 2014). 2000 synchronized L4 worms were collected into organic solvent resistant Eppendorf tubes and suspended in 250μL precooled methanol. Subsequently, 875 μL MTBE was added. Worms were lysed with ice cold ultrasonic bath for 30 minutes. Phase separation was induced by adding 210 μL water followed by sonication for 15 minutes. The organic supernatant was collected after centrifugation at 4°C. Re-extraction of the lower phase was done by adding additional 325 μL MTBE. The organic supernatant was collected and combined with those collected previously and evaporated with under a stream of nitrogen at room temperature and then stored at −80°C. Before analysis, lipids were dissolved in 200 μL acetonitrile / isopropanol / water (65/30/5, v/v/v) and a 100 μL aliquot was used for LC-MS analysis per sample.

### Liquid chromatography

Lipid separation was done with the Bruker Elute UPLC system (Brucker) on a Bruker intensity Solo 2 C18 column (100 x 2.1 mm ID, 1.7 μm particle size) (Brucker). The column temperature was held at 55°C and the lipids were separated with a gradient from eluent A (acetonitrile / water (60:40, 10 mM NH4 Formate, 0.1% formic acid (v/v)) to eluent B (isopropanol / acetonitrile (90:10, 10 mM NH4 Formate, 0.1% formic acid (v/v)) with a flow rate of 400 μL/min. Gradient conditions: After an isocratic step of 40% B for 2 min, then 43% B for 0.1 min, 50% B for 10min, 54% B for 0.1min, 70% B for 6min and 99% B for 0.1 min. After returning to initial conditions, the column was re-equilibrated for 2 min with the starting conditions.

### Trapped ion mobility-PASEF mass spectrometry

Output from liquid chromatography was injected into a trapped ion mobility-quadrupole time-of-flight mass spectrometer (timsTOF Pro MS) (Brucker) through an electrospray ion source. The mass spectrometry was powered by the Parallel Accumulation Serial Fragmentation (PASEF). Analysis was performed twice for each sample in both positive and negative electrospray modes. The parameter was modified based on a published method (Vasilopoulou et al., 2020). The mass spectra were scanned over the range of m/z 100 to m/z 1350 while the ion mobility was recorded from 0.55 Vs cm^2^ to 1.90 Vs cm^2^. Precursors within ± 1 Th were isolated for MS/MS acquisition. 10 eV was used as collision energy for fragmentation. The acquisition cycle was composed of one full TIMS-MS scan and 2 PASEF MS/MS scans, taking 0.31s. Precursor ions with an intensity within the range of 100-4000 counts was rescheduled. PASEF exclusion release Time was set to 0.1 min. IMS ion charge control was 5×10^6^. The calibration of TIMS was done with the Agilent ESI LC/MS tuning mix (Agilent Technology). The selected ions for the calibration of positive modes were: [m/z, 1/K0: (322.0481, 0.7318 Vs cm^−2^), (622.0289,0.9848 Vs cm^−2^), (922.0097, 1.1895 Vs cm^−2^), (1221,9906, 1.3820 Vs cm^−2^)] while those for negative modes were: [m/z, 1/K0: (601.9790, 0.8782 Vs cm^−2^), (1033.9882, 1.2526 Vs cm^−2^), (1333.9690, 1.4016 Vs cm^−2^), (1633.9498, 1.5731 Vs cm^−2^)]. For negative mode, only mass was calibrated but not the mobility.

### Lipid annotation and statistical analysis

The mass spectrometry data was analyzed with MetaboScape version 5.0 (Bruker Daltonics), which extracts retention time, m/z, collision cross section and intensity for each chemical detected while assigning MS/MS spectra to it. After obtaining the merged bucket table from both the positive and negative modes, detected chemicals were firstly annotated with Spectral Libraries: MSDIAL-Tandem Mass Spectral Atlas-VS68-pos and MSDIAL-Tandem Mass Spectral Atlas-VS68-neg. and then annotated with analyte lists of fatty aldehydes [FA06], fatty alcohols [FA05], and fatty Acids and Conjugates [FA01], sphingolipids [SP] from LIPID MAPS (https://www.lipidmaps.org/). Finally, entries in the bucket table were annotated with SmartFormula. The intensity was normalized with probabilistic quotient normalization (Dieterle et al., 2006). The following lipid categories were searched: Triacylglycerol (TAG), Phosphatidylethanolamine (PE), Phosphatidylcholine (PC), Phosphatidylserine (PS), Phosphoinositide (PI), ether linked phospholipid, sphingolipid, sphingomyelin, fatty aldehyde/fatty acid (MS information was insufficient to distinguish fatty aldehyde and fatty alcohol), fatty acids and diacylglycerol (DAG). The average intensity of the two technical replicates (0 value was considered as missing value (NA) and therefore ignored) for each chemical detected in that category were summed up to get the total amount for the corresponding categories.

### Statistical analysis

Data were shown as mean ± SEM and Student’s t-test (unpaired, two-tailed) in GraphPad Prism or Microsoft Excel was applied for statistical analysis unless otherwise indicated.

## Acknowledgments

We thank Meng Wang and King L. Chow for RNAi reagents and protocols. Jihong Bai, Christian Frøkjær-Jensen and Bob Goldstein for Mos1 and CRISPR reagents. Pui Shuen Wong at the HKUST Bioscience Central Research Facility for RNA sequencing and lipidomic analysis. Yan Li and Xingyu Yang for technical advice. Some strains were provided by the CGC, which is funded by NIH Office of Research Infrastructure Programs (P40 OD010440). The work was supported by RGC GRF 16102118 to HYM.

---

**Figure S1.**
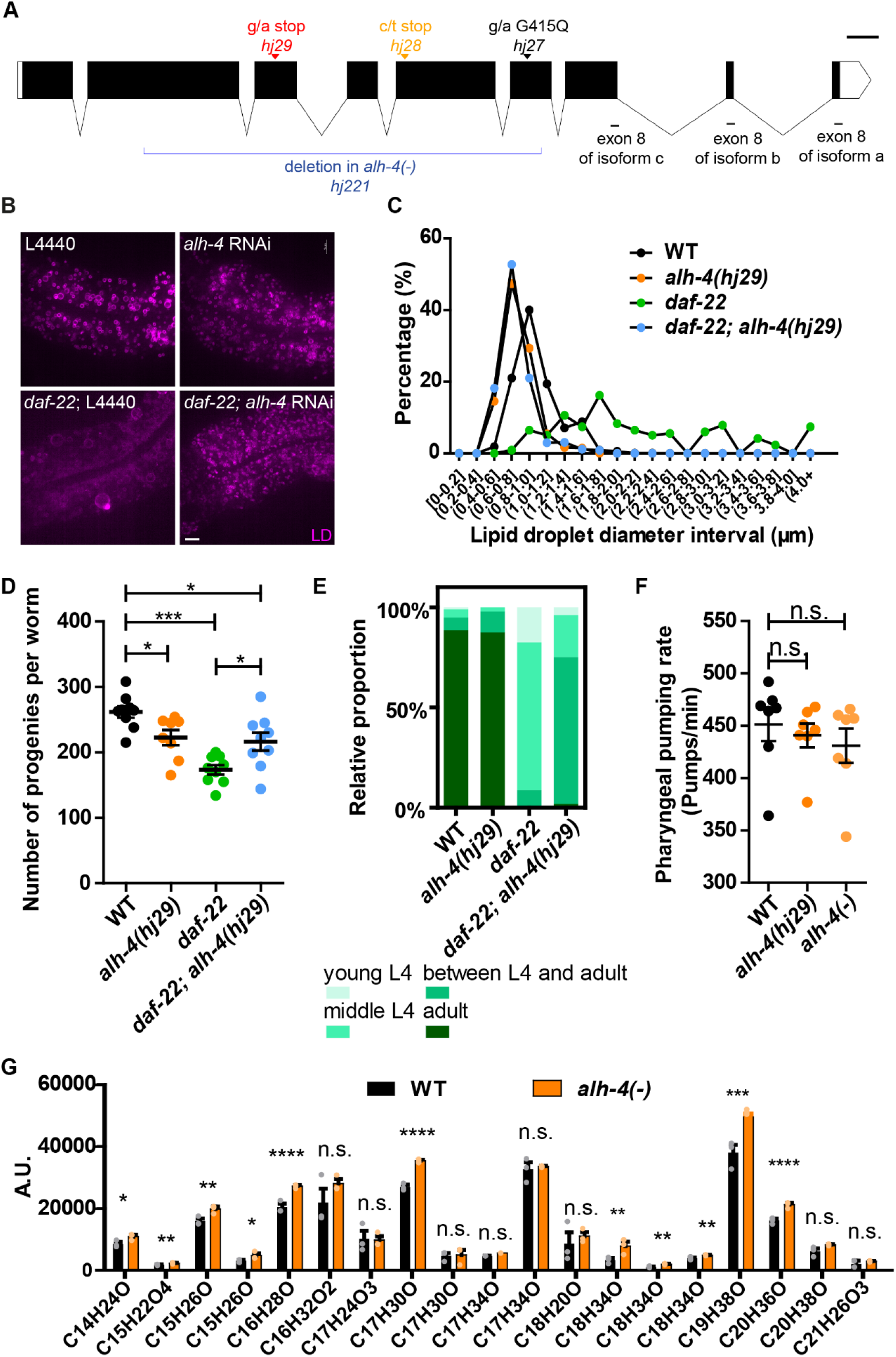
Phenotypic characterization of *alh-4* mutant worms. (A) Schematic representation of the *alh-4* gene structure. Black boxes, exons; white boxes, untranslated region; line, splicing. The mutation sites in *hj28* and *hj29* are indicated by red triangle and yellow triangle, respectively. The single nucleotide substitution in both alleles result in a premature stop codon. The mutation site in *hj27* is indicated by black triangle and the single nucleotide substitution results in a missense mutation Gly 415 Glu. The deletion region in *alh-4* in *hj221* is indicated with the blue line. *alh-4(hj221)* is labeled as *alh-4(−)* in all figures. Scale bar = 100bp. (B) Representative images of lipid droplets in the second intestinal segment of L4 worms. Lipid droplets were labeled with mRuby::DGAT-2 (*hjSi112*). Scale bar = 5μm. (C) Frequency distribution of LD diameter of wild type (WT) and mutant worms of indicated genotypes. Total number of the lipid droplet measured: WT = 1055, *alh-4(hj29)* = 1628, *daf-22(ok693)* = 216, *alh-4(hj29); daf-22(ok693)* = 943. (D) Quantification of number of progenies of wild type (WT) and mutant worms of indicated genotypes. (E) Percentage of 50 synchronized worms in the young L4, middle L4, between L4 and adult or adult developmental stage observed at 70 hours after egg laying. (F) Quantification of the pharyngeal pumping rate of WT, *alh-4(hj29)* and *alh-4(hj221)* worms. n = 7 for each group. (G) Quantification of indicated fatty aldehydes and fatty alcohols in wild type (WT) and *alh-4(−)* worms with LC-MS. A significant increase in multiple species of fatty aldehyde/fatty alcohol was observed in *alh-4(−)* worms in comparison with WT worms. n = 3 for each group. Each dot represents the average value of two technical replicates of each biological sample. In the plot, mean ± SEM of each group is shown. For all statistical analysis, *p < 0.05, **p < 0.01, ***p < 0.001; ****p < 0.0001 (unpaired Student’s t-test).

**Figure S2.**
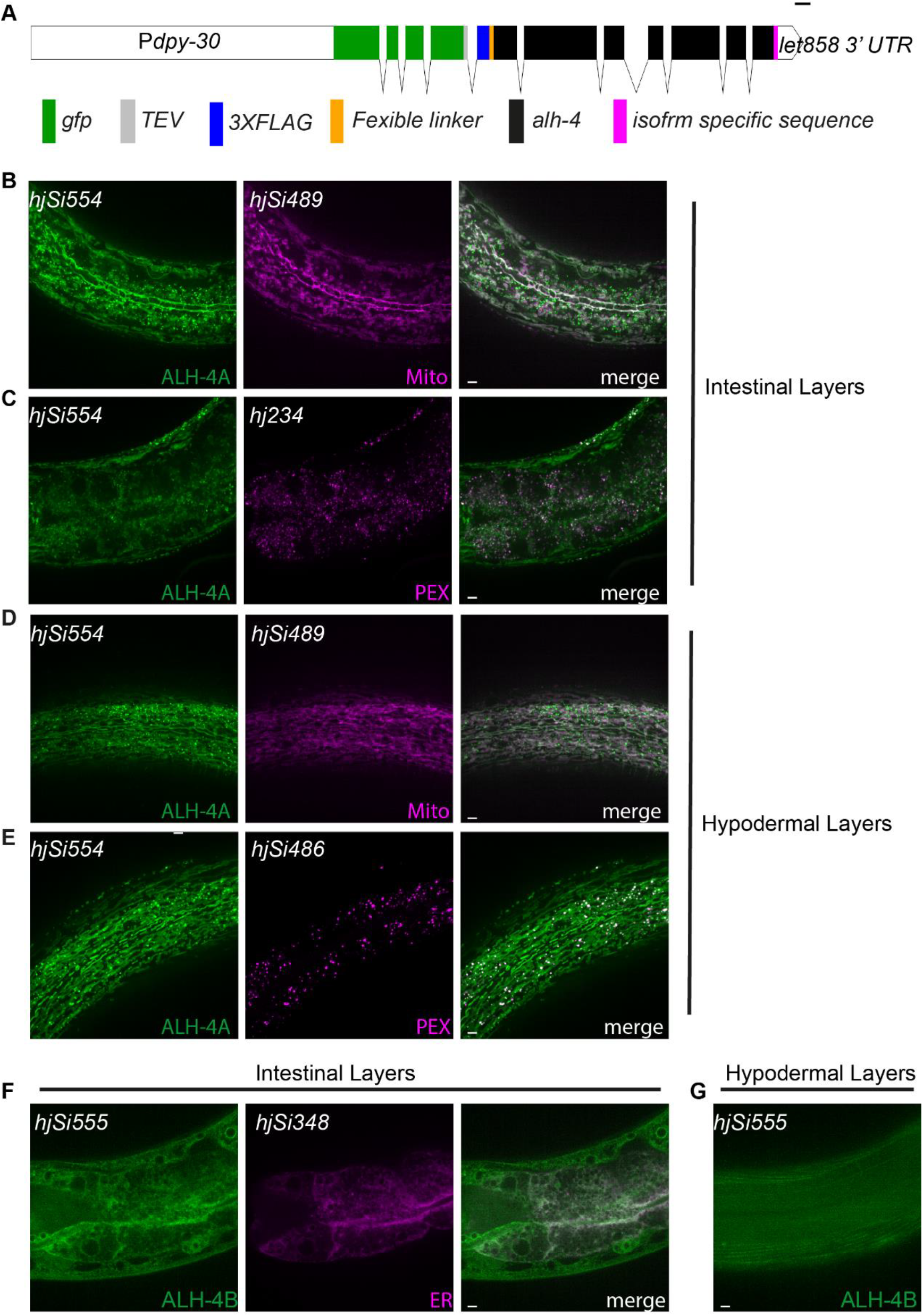
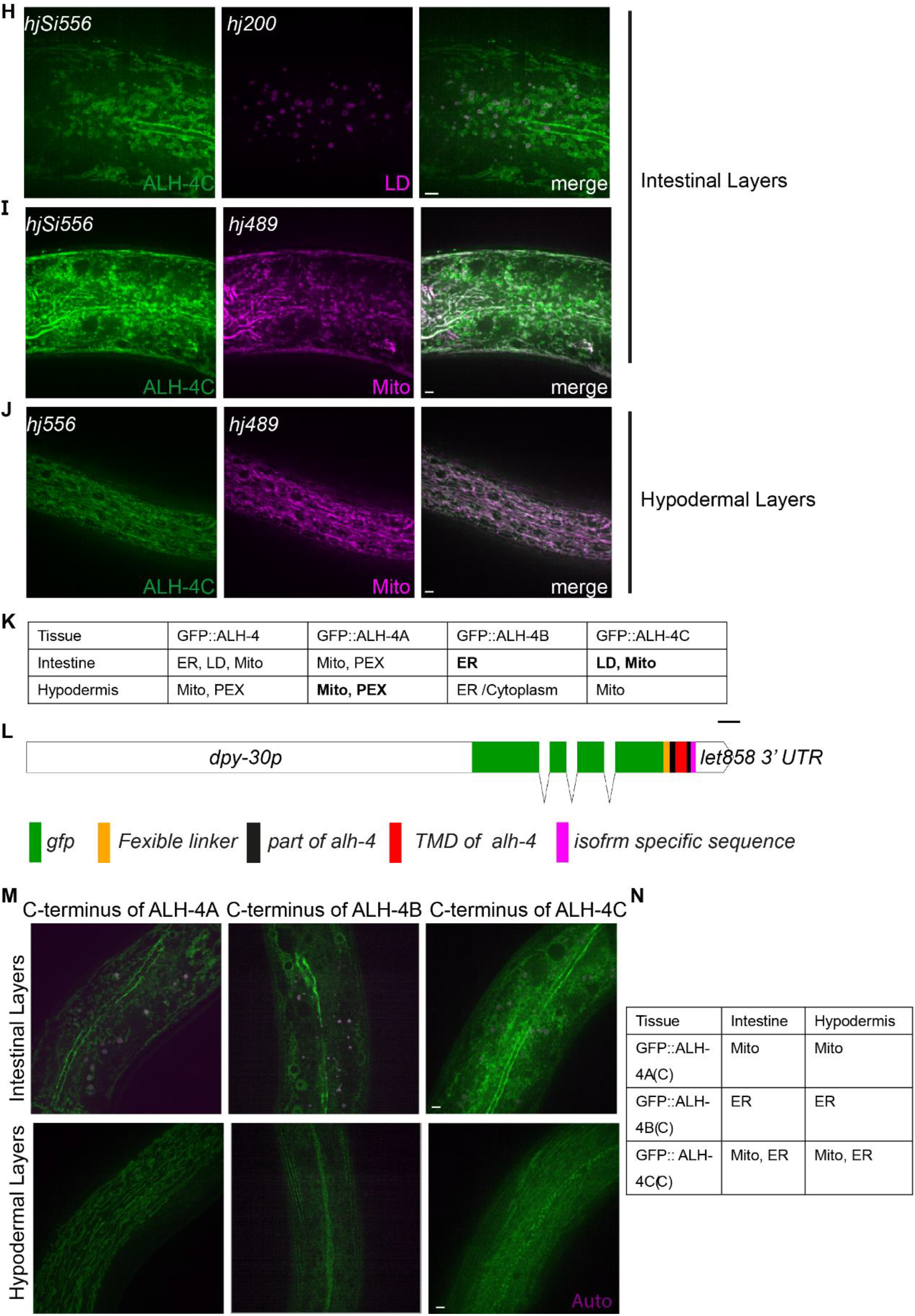
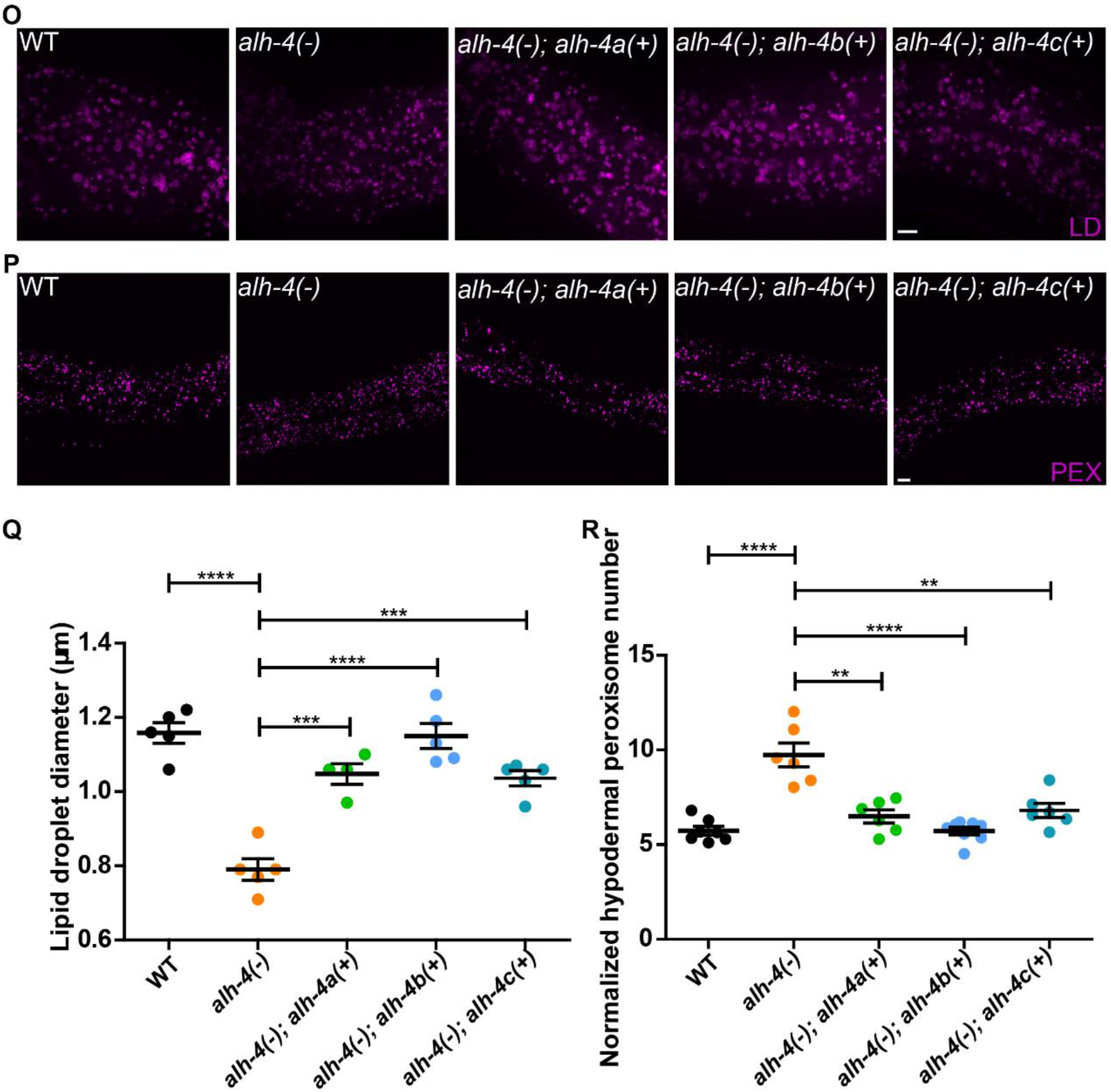
Subcellular localization of ALH-4 and transgenic rescue of *alh-4(−)* worms. (A) Schematic representation of the structure of transgenes expressing GFP tagged ALH-4 isoforms. The magenta box indicates isoform specific sequence. ALH-4A isoform: AAATATGCTCGCAATCTTCATTGA encoding KYARNLH* ALH-4B isoform: TTCATTTTCCGCTTCTCGGCATAA encoding FIFRFSA*. ALH-4C isoform: GTATGCAGAGGAAAATCAGCTCAATAG encoding VCRGKSAQ*. Scale bar = 100bp. (B - E) GFP::ALH-4A (*hjSi554*) colocalized with mitochondria labeled with TOMM-20N::mRuby (*hjSi489*) and peroxisomes labeled with tagRFP::DAF-22 (*hj234*) in the intestine and tagRFP::PTS1 (*hjSi486*) in the hypodermis. (F) GFP::ALH-4B (*hjSi555*) colocalized with the ER labeled with ACS-22::tagRFP (*hjSi348*) in the intestine. (G) No distinct pattern of GFP::ALH-4B was detected in the hypodermis. (H - I) In the intestine, GFP::ALH-4C (*hjSi556*) colocalized with lipid droplets labeled with DHS-3::mRuby (*hj200*) and mitochondria labeled with TOMM-20N::mRuby (*hjSi489*). (J) In the hypodermis, GFP::ALH-4C colocalized with mitochondria labeled with TOMM-20N::mRuby (*hjSi489*). (K) Table summarizing the subcellular localization of different ALH-4 isoforms in the intestine and hypodermis. (L) Schematic representation of the structure of transgenes expressing GFP fused with the C-terminus of ALH-4 isoforms. Scale bar = 100bp. (M) Representative images showing the subcellular localization of GFP fused with C-terminus of ALH-4 isoforms in the intestine (top) and in the hypodermis (bottom). (N) Table summarizing the results shown in (M). (O) Transgenic rescue of *alh-4(−)* worms with specific ALH-4 isoforms. Representative images showing intestinal lipid droplets labeled with DHS-3::mRuby(*hj200*). (P) As in (O), but with representative images showing the hypodermal peroxisomes labeled with tagRFP::PTS1(*hjSi486*). (Q) Quantification of intestinal lipid droplet diameter. n = 5 for each group. (R) Quantification the hypodermal peroxisome number normalized with the length of the worm in the field of view. n > 6 for each group. In each plot, mean ± SEM of each group is shown. *p < 0.05, **p < 0.01, ***p < 0.001; ****p < 0.0001 (unpaired Student’s t-test). PEX, peroxisome; LD, lipid droplet. Scale bar = 5μm.

**Figure S3.**
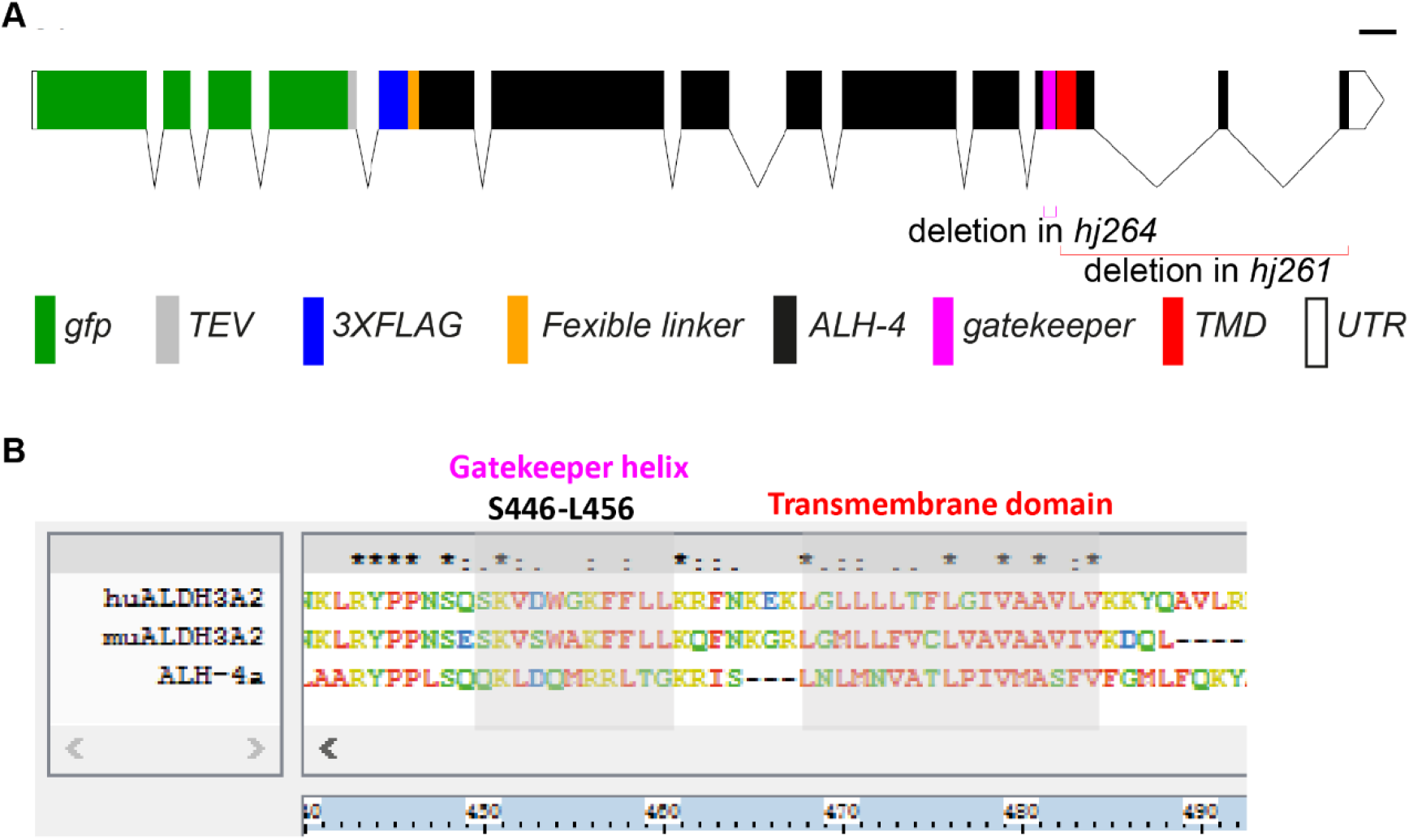
Engineered mutations for the disruption of the membrane anchor or gate-keeper region of ALH-4. (A) Schematic representation of *gfp::alh-4*(*Δgk*) (*hj264*) and *gfp::alh-4*(*ΔC*) (*hj261*), which had the gate-keeper region and C-terminal membrane anchor deleted, respectively. The deleted region is indicated. TMD, transmembrane domain. Scale bar = 100bp. (B) Sequence alignment of human and mouse ALDH3A2 and *C. elegans* ALH-4A (with Clustal X2). Residue color scheme: hydrophobic (A, L, M, P, I, F, W, V), red; polar (N, C, Q, T, Y, G, S), green; negative charged (D, E), blue; positive charged (H, R, K), yellow. Alignment quality display: ‘*’ for a single, fully conserved residue; ‘:’ for fully conserved of some “strong groups”; ‘.’ for fully conserved of some “weak groups”; Corresponding regions are in grey areas.

**Figure S4.**
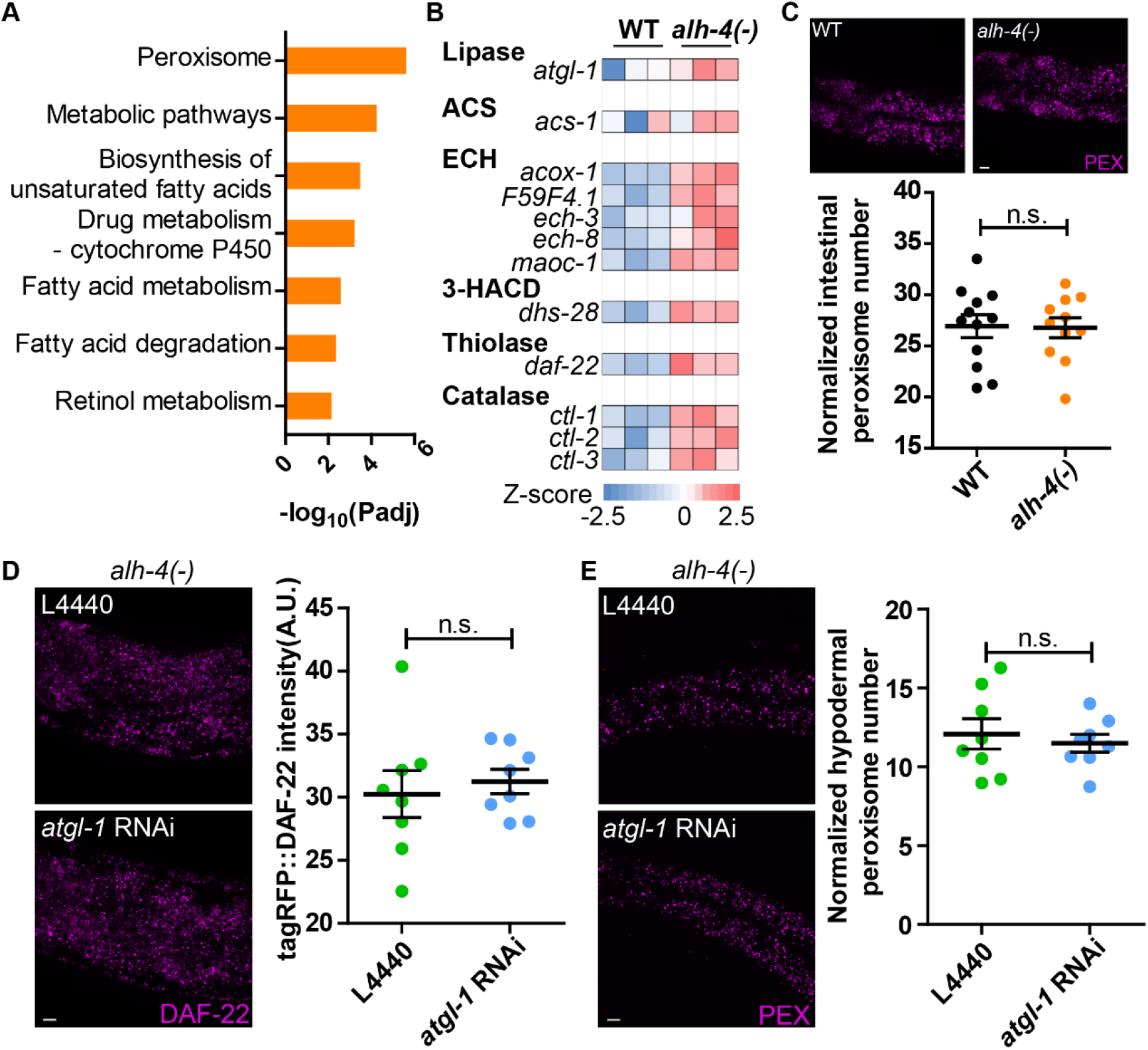
Transcriptomic analysis of *alh-4(−)* worms. (A) KEGG pathway analysis of the differentially expressed genes (fold change > 2 and p-value < 0.05) between wild type (WT) and *alh-4(−)* worms. The analysis was performed with https://biit.cs.ut.ee/gprofiler/gost. The peroxisome pathway and additional pathways related to lipid metabolism were returned as top hits. (B) Heatmap showing the differential expression of genes encoding peroxisomal fatty acid β-oxidation enzymes in WT and *alh-4* worms. The FPKM value is converted to Z-scores. High expression is indicated with red, while low expression is indicated with blue. n = 3 for each group. Only genes showing significant difference (p-value < 0.1) in the expression are shown except *acs-1* (p-value = 0.30). (C) Representative images (Z-stack, 0.5 μm intervals, 10 slices) showing the intestinal peroxisomes labeled with mRuby::PTS1 (*hjSi548*) (top). Quantification of intestinal peroxisome number (bottom). n =11 each group. (D) Representative images showing tagRFP::DAF-22 (*hj234*) in *alh-4(−)* worms that were subject to control (L4440) or *atgl-1* RNAi (top). Quantification of tagRFP::DAF-22 fluorescence. n = 8 for each group. (E) As in (D), but with hypodermal peroxisomes labeled with tagRFP::PTS1(*hjsi486*) (top). Quantification of peroxisome number normalized with length of the worms in the field (bottom). n = 8 for each group. In each plot, mean ± SEM of each group is shown. *p < 0.05, **p < 0.01, ***p < 0.001; ****p < 0.0001 (unpaired Student’s t-test). PEX, peroxisome. Scale bar = 5μm.

**Figure S5.**
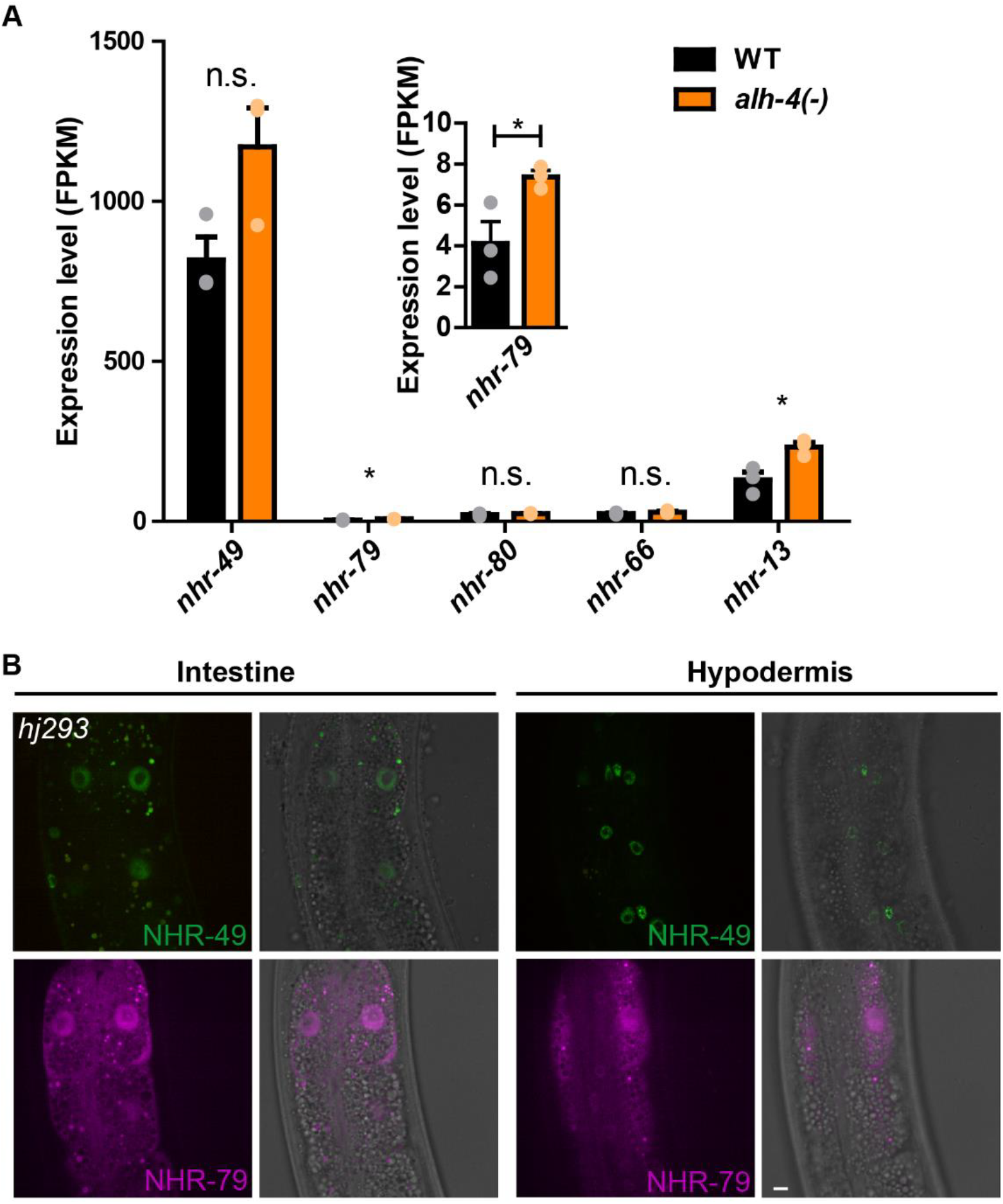
The expression level and subcellular localization of NHR-49 and NHR-79. (A) Histogram showing the mRNA levels of *nhr-49, nhr-79, nhr-80, nhr-66* and *nhr-13*, from RNA sequencing of wild type (WT) and *alh-4(−)* worms. n = 3 for each group. In the plot, mean ± SEM of each group is shown. For all statistical analysis, *p < 0.05, **p < 0.01, ***p < 0.001, ****p < 0.0001, n.s. not significant (unpaired Student’s t-test). (B) Visualization of NHR-49::GFP (*hj*293) from the endogenous locus (top) and the visualization of mRuby::NHR-79 (*hjEx26[nhr-79p::nhr-79]*(bottom). Both NHR-49::GFP and mRuby::NHR-79 could be detected in the nuclei of intestinal and hypodermal cells. Scale bar = 5μm.

**Figure S6.**
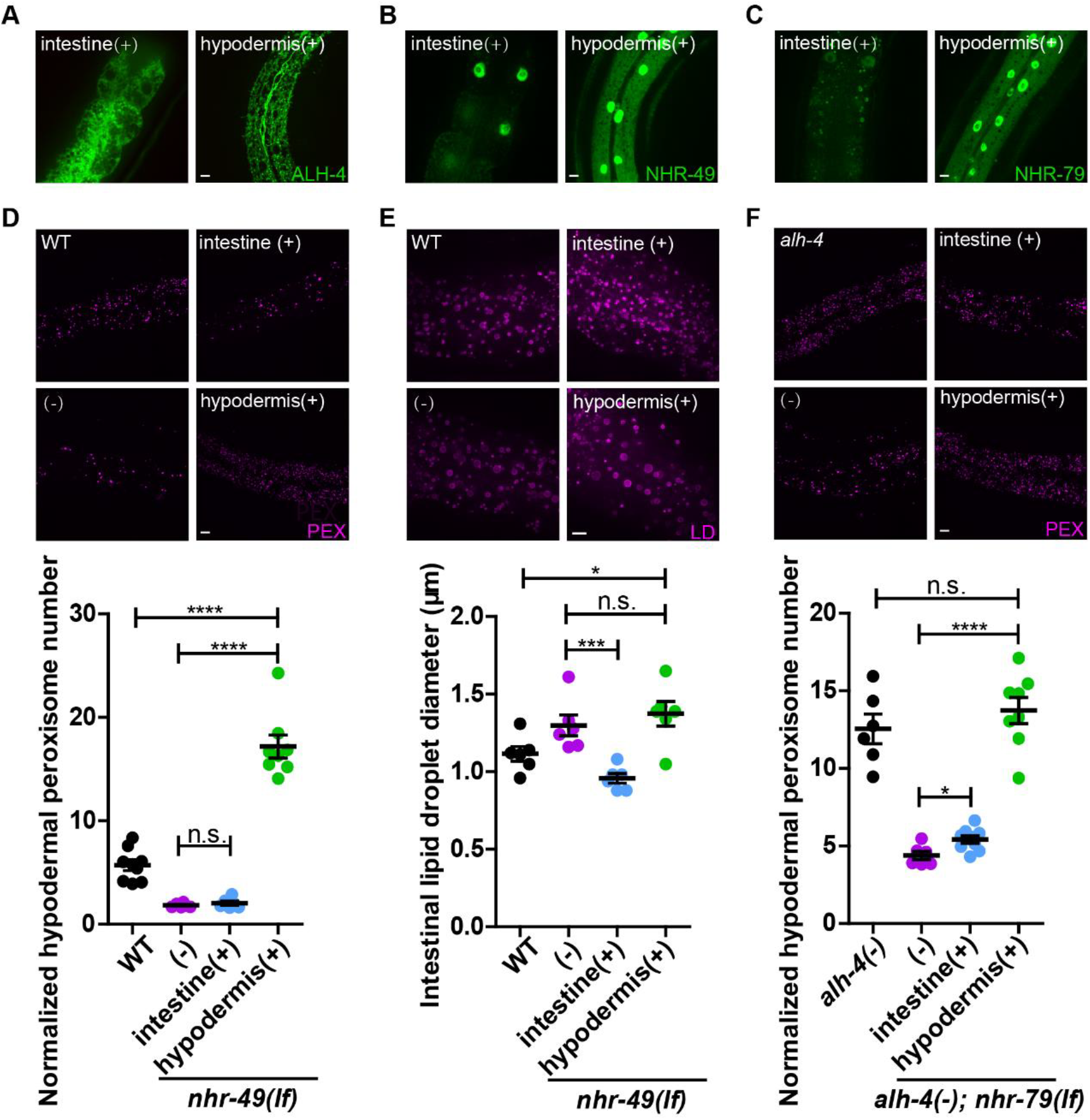
Tissue specific rescue of *alh-4(−)*, *nhr-49(lf)* and *nhr-79(lf)* worms. (A) Representative images showing the intestinal specific expression of GFP::ALH-4 in worms carrying *hjSi501*(left) and the hypodermal specific expression of GFP::ALH-4 in worms *hjSi500* (right). (B) Representative images showing the intestinal specific expression of NHR-49A/B::GFP in worms carrying *hjSi531* (left) and the hypodermal specific expression of NHR-49A/B::GFP in worms carrying *hjSi537* (right). (C) Representative images showing the intestinal specific expression of GFP::NHR-79A in worms carrying *hjSi553*(left) and the hypodermal specific expression of GFP::NHR-79A in worms carrying *hjSi539* (right). (D) Representative images showing hypodermal peroxisomes labeled with tagRFP::PTS1 (*hjSi486)* (top). Quantification of hypodermal peroxisome number in wild type (WT) and mutant worms of indicated genotypes. n > 6 for each group. (E) As in (D), except with intestinal lipid droplets labeled with DHS-3::mRuby (*hj200)* (top). Quantification of lipid droplets in the second intestinal segment (bottom). n = 6 for each group. (F) As in (D), except with WT and *alh-4(−)* and *nhr-79(−)* mutant worms of indicated genotypes. Quantification of hypodermal peroxisome number (bottom). n > 6 for each group. In each plot, mean ± SEM of each group is shown. For all statistical analysis, *p < 0.05, **p < 0.01, ***p < 0.001; ****p < 0.0001, n.s. not significant. PEX, peroxisome; LD, lipid droplet. Scale bar = 5μm.

## References

An, J.H., and Blackwell, T.K. (2003). SKN-1 links C. elegans mesendodermal specification to a conserved oxidative stress response. Genes Dev. 17, 1882–1893, doi:10.1101/gad.1107803.

Angelo, G., and Van Gilst, M.R. (2009). Starvation protects germline stem cells and extends reproductive longevity in C. elegans. Science 326, 954–958, doi:10.1126/science.1178343.

Arda, H.E., Taubert, S., MacNeil, L.T., Conine, C.C., Tsuda, B., Van Gilst, M., Sequerra, R., Doucette-Stamm, L., Yamamoto, K.R., and Walhout, A.J.M. (2010). Functional modularity of nuclear hormone receptors in a Caenorhabditis elegans metabolic gene regulatory network. Mol Syst Biol 6, 367, doi:10.1038/msb.2010.23.

Ashibe, B., and Motojima, K. (2009). Fatty aldehyde dehydrogenase is up-regulated by polyunsaturated fatty acid via peroxisome proliferator-activated receptor alpha and suppresses polyunsaturated fatty acid-induced endoplasmic reticulum stress. FEBS J 276, 6956–6970, doi:10.1111/j.1742-4658.2009.07404.x.

Ashibe, B., Hirai, T., Higashi, K., Sekimizu, K., and Motojima, K. (2007). Dual Subcellular Localization in the Endoplasmic Reticulum and Peroxisomes and a Vital Role in Protecting against Oxidative Stress of Fatty Aldehyde Dehydrogenase Are Achieved by Alternative Splicing. J. Biol. Chem. 282, 20763–20773, doi:10.1074/jbc.M611853200.

Bailey, A.P., Koster, G., Guillermier, C., Hirst, E.M.A., MacRae, J.I., Lechene, C.P., Postle, A.D., and Gould, A.P. (2015). Antioxidant Role for Lipid Droplets in a Stem Cell Niche of Drosophila. Cell 163, 340–353, doi:10.1016/j.cell.2015.09.020.

Brock, T.J., Browse, J., and Watts, J.L. (2006). Genetic regulation of unsaturated fatty acid composition in C. elegans. PLoS Genet. 2, e108, doi:10.1371/journal.pgen.0020108.

Burkewitz, K., Morantte, I., Weir, H.J.M., Yeo, R., Zhang, Y., Huynh, F.K., Ilkayeva, O.R., Hirschey, M.D., Grant, A.R., and Mair, W.B. (2015). Neuronal CRTC-1 governs systemic mitochondrial metabolism and lifespan via a catecholamine signal. Cell 160, 842–855, doi:10.1016/j.cell.2015.02.004.

Cao, Z., Hao, Y., Fung, C.W., Lee, Y.Y., Wang, P., Li, X., Xie, K., Lam, W.J., Qiu, Y., Tang, B.Z., et al. (2019). Dietary fatty acids promote lipid droplet diversity through seipin enrichment in an ER subdomain. Nat Commun 10, 2902, doi:10.1038/s41467-019-10835-4.

Chung, H.-L., Wangler, M.F., Marcogliese, P.C., Jo, J., Ravenscroft, T.A., Zuo, Z., Duraine, L., Sadeghzadeh, S., Li-Kroeger, D., Schmidt, R.E., et al. (2020). Loss- or Gain-of-Function Mutations in ACOX1 Cause Axonal Loss via Different Mechanisms. Neuron 106, 589–606.e6, doi:10.1016/j.neuron.2020.02.021.

Costello, J.L., Castro, I.G., Camões, F., Schrader, T.A., McNeall, D., Yang, J., Giannopoulou, E.-A., Gomes, S., Pogenberg, V., Bonekamp, N.A., et al. (2017). Predicting the targeting of tail-anchored proteins to subcellular compartments in mammalian cells. J. Cell. Sci. 130, 1675–1687, doi:10.1242/jcs.200204.

Davis, M.W., Hammarlund, M., Harrach, T., Hullett, P., Olsen, S., and Jorgensen, E.M. (2005). Rapid single nucleotide polymorphism mapping in C. elegans. BMC Genomics 6, 118, doi:10.1186/1471-2164-6-118.

De Laurenzi, V., Rogers, G.R., Hamrock, D.J., Marekov, L.N., Steinert, P.M., Compton, J.G., Markova, N., and Rizzo, W.B. (1996). Sjögren-Larsson syndrome is caused by mutations in the fatty aldehyde dehydrogenase gene. Nat Genet 12, 52–57, doi:10.1038/ng0196-52.

Demozay, D., Mas, J.-C., Rocchi, S., and Van Obberghen, E. (2008). FALDH reverses the deleterious action of oxidative stress induced by lipid peroxidation product 4-hydroxynonenal on insulin signaling in 3T3-L1 adipocytes. Diabetes 57, 1216–1226, doi:10.2337/db07-0389.

Dickinson, D.J., Ward, J.D., Reiner, D.J., and Goldstein, B. (2013). Engineering the Caenorhabditis elegans genome using Cas9-triggered homologous recombination. Nat. Methods 10, 1028–1034, doi:10.1038/nmeth.2641.

Dickinson, D.J., Pani, A.M., Heppert, J.K., Higgins, C.D., and Goldstein, B. (2015). Streamlined genome engineering with a self-excising drug selection cassette. Genetics 200, 1035–1049, doi:10.1534/genetics.115.178335.

Dieterle, F., Ross, A., Schlotterbeck, G., and Senn, H. (2006). Probabilistic quotient normalization as robust method to account for dilution of complex biological mixtures. Application in 1H NMR metabonomics. Anal Chem 78, 4281–4290, doi:10.1021/ac051632c.

Dubois, V., Eeckhoute, J., Lefebvre, P., and Staels, B. (2017). Distinct but complementary contributions of PPAR isotypes to energy homeostasis. J Clin Invest 127, 1202–1214, doi:10.1172/JCI88894.

Ebenezer, D.L., Fu, P., Ramchandran, R., Ha, A.W., Putherickal, V., Sudhadevi, T., Harijith, A., Schumacher, F., Kleuser, B., and Natarajan, V. (2020). S1P and plasmalogen derived fatty aldehydes in cellular signaling and functions. Biochim Biophys Acta Mol Cell Biol Lipids 1865, 158681, doi:10.1016/j.bbalip.2020.158681.

Evans, R.M., and Mangelsdorf, D.J. (2014). Nuclear Receptors, RXR, and the Big Bang. Cell 157, 255–266, doi:10.1016/j.cell.2014.03.012.

Fidaleo, M., Fanelli, F., Ceru, M.P., and Moreno, S. (2014). Neuroprotective properties of peroxisome proliferator-activated receptor alpha (PPARα) and its lipid ligands. Curr Med Chem 21, 2803–2821, doi:10.2174/0929867321666140303143455.

Folick, A., Oakley, H.D., Yu, Y., Armstrong, E.H., Kumari, M., Sanor, L., Moore, D.D., Ortlund, E.A., Zechner, R., and Wang, M.C. (2015). Aging. Lysosomal signaling molecules regulate longevity in Caenorhabditis elegans. Science 347, 83–86, doi:10.1126/science.1258857.

Fredens, J., and Færgeman, N.J. (2012). Quantitative proteomics by amino acid labeling identifies novel NHR-49 regulated proteins in C. elegans. Worm 1, 66–71, doi:10.4161/worm.19044.

Frøkjær-Jensen, C., Davis, M.W., Ailion, M., and Jorgensen, E.M. (2012). Improved Mos1-mediated transgenesis in C. elegans. Nat. Methods 9, 117–118, doi:10.1038/nmeth.1865.

Fu, J., Gaetani, S., Oveisi, F., Lo Verme, J., Serrano, A., Rodríguez De Fonseca, F., Rosengarth, A., Luecke, H., Di Giacomo, B., Tarzia, G., et al. (2003). Oleylethanolamide regulates feeding and body weight through activation of the nuclear receptor PPAR-alpha. Nature 425, 90–93, doi:10.1038/nature01921.

Gloerich, J., Ijlst, L., Wanders, R.J.A., and Ferdinandusse, S. (2006). Bezafibrate induces FALDH in human fibroblasts; implications for Sjögren-Larsson syndrome. Mol Genet Metab 89, 111–115, doi:10.1016/j.ymgme.2006.05.009.

Goh, G.Y.S., Winter, J.J., Bhanshali, F., Doering, K.R.S., Lai, R., Lee, K., Veal, E.A., and Taubert, S. (2018). NHR-49/HNF4 integrates regulation of fatty acid metabolism with a protective transcriptional response to oxidative stress and fasting. Aging Cell 17, e12743, doi:10.1111/acel.12743.

Guéraud, F., Atalay, M., Bresgen, N., Cipak, A., Eckl, P.M., Huc, L., Jouanin, I., Siems, W., and Uchida, K. (2010). Chemistry and biochemistry of lipid peroxidation products. Free Radic Res 44, 1098–1124, doi:10.3109/10715762.2010.498477.

Hayes, G.D., Frand, A.R., and Ruvkun, G. (2006). The mir-84 and let-7 paralogous microRNA genes of Caenorhabditis elegans direct the cessation of molting via the conserved nuclear hormone receptors NHR-23 and NHR-25. Development 133, 4631–4641, doi:10.1242/dev.02655.

James, P.F., and Zoeller, R.A. (1997). Isolation of animal cell mutants defective in long-chain fatty aldehyde dehydrogenase. Sensitivity to fatty aldehydes and Schiff’s base modification of phospholipids: implications for Sj-ogren-Larsson syndrome. J. Biol. Chem. 272, 23532–23539, doi:10.1074/jbc.272.38.23532.

Jeninga, E.H., Gurnell, M., and Kalkhoven, E. (2009). Functional implications of genetic variation in human PPARgamma. Trends Endocrinol Metab 20, 380–387, doi:10.1016/j.tem.2009.04.005.

Kanetake, T., Sassa, T., Nojiri, K., Sawai, M., Hattori, S., Miyakawa, T., Kitamura, T., and Kihara, A. (2019). Neural symptoms in a gene knockout mouse model of Sjögren-Larsson syndrome are associated with a decrease in 2-hydroxygalactosylceramide. FASEB J 33, 928–941, doi:10.1096/fj.201800291R.

Keller, M.A., Zander, U., Fuchs, J.E., Kreutz, C., Watschinger, K., Mueller, T., Golderer, G., Liedl, K.R., Ralser, M., Kräutler, B., et al. (2014). A gatekeeper helix determines the substrate specificity of Sjögren-Larsson Syndrome enzyme fatty aldehyde dehydrogenase. Nat Commun 5, 4439, doi:10.1038/ncomms5439.

Klemm, R.W., Norton, J.P., Cole, R.A., Li, C.S., Park, S.H., Crane, M.M., Li, L., Jin, D., Boye-Doe, A., Liu, T.Y., et al. (2013). A conserved role for atlastin GTPases in regulating lipid droplet size. Cell Rep 3, 1465–1475, doi:10.1016/j.celrep.2013.04.015.

Kliewer, S.A., Umesono, K., Noonan, D.J., Heyman, R.A., and Evans, R.M. (1992). Convergence of 9-cis retinoic acid and peroxisome proliferator signalling pathways through heterodimer formation of their receptors. Nature 358, 771–774, doi:10.1038/358771a0.

Kojetin, D.J., Matta-Camacho, E., Hughes, T.S., Srinivasan, S., Nwachukwu, J.C., Cavett, V., Nowak, J., Chalmers, M.J., Marciano, D.P., Kamenecka, T.M., et al. (2015). Structural mechanism for signal transduction in RXR nuclear receptor heterodimers. Nat Commun 6, 8013, doi:10.1038/ncomms9013.

Lehtinen, M.K., Yuan, Z., Boag, P.R., Yang, Y., Villén, J., Becker, E.B.E., DiBacco, S., de la Iglesia, N., Gygi, S., Blackwell, T.K., et al. (2006). A conserved MST-FOXO signaling pathway mediates oxidative-stress responses and extends life span. Cell 125, 987–1001, doi:10.1016/j.cell.2006.03.046.

Li, X., Lam, W.J., Cao, Z., Hao, Y., Sun, Q., He, S., Mak, H.Y., and Qu, J.Y. (2015). Integrated femtosecond stimulated Raman scattering and two-photon fluorescence imaging of subcellular lipid and vesicular structures. J Biomed Opt 20, 110501, doi:10.1117/1.JBO.20.11.110501.

Liang, B., Ferguson, K., Kadyk, L., and Watts, J.L. (2010). The role of nuclear receptor NHR-64 in fat storage regulation in Caenorhabditis elegans. PLoS ONE 5, e9869, doi:10.1371/journal.pone.0009869.

Lin, C.-C.J., and Wang, M.C. (2017). Microbial metabolites regulate host lipid metabolism through NR5A-Hedgehog signalling. Nat. Cell Biol. 19, 550–557, doi:10.1038/ncb3515.

Liu, L., MacKenzie, K.R., Putluri, N., Maletić-Savatić, M., and Bellen, H.J. (2017). The Glia-Neuron Lactate Shuttle and Elevated ROS Promote Lipid Synthesis in Neurons and Lipid Droplet Accumulation in Glia via APOE/D. Cell Metab 26, 719–737.e6, doi:10.1016/j.cmet.2017.08.024.

Magner, D.B., Wollam, J., Shen, Y., Hoppe, C., Li, D., Latza, C., Rottiers, V., Hutter, H., and Antebi, A. (2013). The NHR-8 nuclear receptor regulates cholesterol and bile acid homeostasis in C. elegans. Cell Metab 18, 212–224, doi:10.1016/j.cmet.2013.07.007.

Mangelsdorf, D.J., Thummel, C., Beato, M., Herrlich, P., Schütz, G., Umesono, K., Blumberg, B., Kastner, P., Mark, M., Chambon, P., et al. (1995). The nuclear receptor superfamily: the second decade. Cell 83, 835–839, doi:10.1016/0092-8674(95)90199-x.

Matyash, V., Liebisch, G., Kurzchalia, T.V., Shevchenko, A., and Schwudke, D. (2008). Lipid extraction by methyl-tert-butyl ether for high-throughput lipidomics. J Lipid Res 49, 1137–1146, doi:10.1194/jlr.D700041-JLR200.

Medina-Rivera, A., Defrance, M., Sand, O., Herrmann, C., Castro-Mondragon, J.A., Delerce, J., Jaeger, S., Blanchet, C., Vincens, P., Caron, C., et al. (2015). RSAT 2015: Regulatory Sequence Analysis Tools. Nucleic Acids Res 43, W50–56, doi:10.1093/nar/gkv362.

Mori, A., Holdorf, A.D., and Walhout, A.J.M. (2017). Many transcription factors contribute to C. elegans growth and fat storage. Genes Cells 22, 770–784, doi:10.1111/gtc.12516.

Motola, D.L., Cummins, C.L., Rottiers, V., Sharma, K.K., Li, T., Li, Y., Suino-Powell, K., Xu, H.E., Auchus, R.J., Antebi, A., et al. (2006). Identification of ligands for DAF-12 that govern dauer formation and reproduction in C. elegans. Cell 124, 1209–1223, doi:10.1016/j.cell.2006.01.037.

Moutinho, M., Codocedo, J.F., Puntambekar, S.S., and Landreth, G.E. (2019). Nuclear Receptors as Therapeutic Targets for Neurodegenerative Diseases: Lost in Translation. Annu Rev Pharmacol Toxicol 59, 237–261, doi:10.1146/annurev-pharmtox-010818-021807.

Mullaney, B.C., Blind, R.D., Lemieux, G.A., Perez, C.L., Elle, I.C., Faergeman, N.J., Van Gilst, M.R., Ingraham, H.A., and Ashrafi, K. (2010). Regulation of C. elegans fat uptake and storage by acyl-CoA synthase-3 is dependent on NR5A family nuclear hormone receptor nhr-25. Cell Metab. 12, 398–410, doi:10.1016/j.cmet.2010.08.013.

Naganuma, T., Takagi, S., Kanetake, T., Kitamura, T., Hattori, S., Miyakawa, T., Sassa, T., and Kihara, A. (2016). Disruption of the Sjögren-Larsson Syndrome Gene Aldh3a2 in Mice Increases Keratinocyte Growth and Retards Skin Barrier Recovery. J Biol Chem 291, 11676–11688, doi:10.1074/jbc.M116.714030.

Nakahara, K., Ohkuni, A., Kitamura, T., Abe, K., Naganuma, T., Ohno, Y., Zoeller, R.A., and Kihara, A. (2012). The Sjögren-Larsson syndrome gene encodes a hexadecenal dehydrogenase of the sphingosine 1-phosphate degradation pathway. Mol. Cell 46, 461–471, doi:10.1016/j.molcel.2012.04.033.

Negre-Salvayre, A., Coatrieux, C., Ingueneau, C., and Salvayre, R. (2008). Advanced lipid peroxidation end products in oxidative damage to proteins. Potential role in diseases and therapeutic prospects for the inhibitors. Br. J. Pharmacol. 153, 6–20, doi:10.1038/sj.bjp.0707395.

Neve, I.A.A., Sowa, J.N., Lin, C.-C.J., Sivaramakrishnan, P., Herman, C., Ye, Y., Han, L., and Wang, M.C. (2020). Escherichia coli Metabolite Profiling Leads to the Development of an RNA Interference Strain for Caenorhabditis elegans. G3 (Bethesda) 10, 189–198, doi:10.1534/g3.119.400741.

Pathare, P.P., Lin, A., Bornfeldt, K.E., Taubert, S., and Van Gilst, M.R. (2012). Coordinate regulation of lipid metabolism by novel nuclear receptor partnerships. PLoS Genet. 8, e1002645, doi:10.1371/journal.pgen.1002645.

Pawlak, M., Lefebvre, P., and Staels, B. (2015). Molecular mechanism of PPARα action and its impact on lipid metabolism, inflammation and fibrosis in non-alcoholic fatty liver disease. J Hepatol 62, 720–733, doi:10.1016/j.jhep.2014.10.039.

Pertea, M., Kim, D., Pertea, G.M., Leek, J.T., and Salzberg, S.L. (2016). Transcript-level expression analysis of RNA-seq experiments with HISAT, StringTie and Ballgown. Nat Protoc 11, 1650–1667, doi:10.1038/nprot.2016.095.

Qi, W., Gutierrez, G.E., Gao, X., Dixon, H., McDonough, J.A., Marini, A.M., and Fisher, L. (2017). The ω-3 fatty acid α-linolenic acid extends Caenorhabditis elegans lifespan via NHR-49/PPARα and oxidation to oxylipins. Aging Cell 16, 1125–1135, doi:10.1111/acel.12651.

Ramachandran, P.V., Mutlu, A.S., and Wang, M.C. (2015). Label-free biomedical imaging of lipids by stimulated Raman scattering microscopy. Curr Protoc Mol Biol 109, 30.3.1–17, doi:10.1002/0471142727.mb3003s109.

Ratnappan, R., Amrit, F.R.G., Chen, S.-W., Gill, H., Holden, K., Ward, J., Yamamoto, K.R., Olsen, C.P., and Ghazi, A. (2014). Germline signals deploy NHR-49 to modulate fatty-acid β-oxidation and desaturation in somatic tissues of C. elegans. PLoS Genet 10, e1004829, doi:10.1371/journal.pgen.1004829.

Reimand, J., Arak, T., Adler, P., Kolberg, L., Reisberg, S., Peterson, H., and Vilo, J. (2016). g:Profiler-a web server for functional interpretation of gene lists (2016 update). Nucleic Acids Res 44, W83–89, doi:10.1093/nar/gkw199.

Rizki, G., Picard, C.L., Pereyra, C., and Lee, S.S. (2012). Host cell factor 1 inhibits SKN-1 to modulate oxidative stress responses in Caenorhabditis elegans. Aging Cell 11, 717–721, doi:10.1111/j.1474-9726.2012.00831.x.

Rizzo, W.B. (2014). Fatty aldehyde and fatty alcohol metabolism: review and importance for epidermal structure and function. Biochim. Biophys. Acta 1841, 377–389, doi:10.1016/j.bbalip.2013.09.001.

Rizzo, W.B., and Craft, D.A. (2000). Sjögren-Larsson syndrome: accumulation of free fatty alcohols in cultured fibroblasts and plasma. J Lipid Res 41, 1077–1081.

Schrader, M., and Fahimi, H.D. (2006). Peroxisomes and oxidative stress. Biochim Biophys Acta 1763, 1755–1766, doi:10.1016/j.bbamcr.2006.09.006.

Sebastian, A., and Contreras-Moreira, B. (2014). footprintDB: a database of transcription factors with annotated cis elements and binding interfaces. Bioinformatics 30, 258–265, doi:10.1093/bioinformatics/btt663.

Serra, L., Chang, D.Z., Macchietto, M., Williams, K., Murad, R., Lu, D., Dillman, A.R., and Mortazavi, A. (2018). Adapting the Smart-seq2 Protocol for Robust Single Worm RNA-seq. Bio Protoc 8, doi:10.21769/BioProtoc.2729.

Singh, S., Brocker, C., Koppaka, V., Chen, Y., Jackson, B.C., Matsumoto, A., Thompson, D.C., and Vasiliou, V. (2013). Aldehyde dehydrogenases in cellular responses to oxidative/electrophilic stress. Free Radical Biology & Medicine 56, 89–101, doi:10.1016/j.freeradbiomed.2012.11.010.

Svensk, E., Ståhlman, M., Andersson, C.-H., Johansson, M., Borén, J., and Pilon, M. (2013). PAQR-2 Regulates Fatty Acid Desaturation during Cold Adaptation in C. elegans. PLoS Genet. 9, e1003801, doi:10.1371/journal.pgen.1003801.

Taubert, S., Ward, J.D., and Yamamoto, K.R. (2011). Nuclear hormone receptors in nematodes: evolution and function. Molecular and Cellular Endocrinology 334, 49–55, doi:10.1016/j.mce.2010.04.021.

Ushio-Fukai, M., and Rehman, J. (2014). Redox and metabolic regulation of stem/progenitor cells and their niche. Antioxid. Redox Signal. 21, 1587–1590, doi:10.1089/ars.2014.5931.

Van Gilst, M.R., Hadjivassiliou, H., Jolly, A., and Yamamoto, K.R. (2005a). Nuclear hormone receptor NHR-49 controls fat consumption and fatty acid composition in C. elegans. PLoS Biol 3, e53, doi:10.1371/journal.pbio.0030053.

Van Gilst, M.R., Hadjivassiliou, H., and Yamamoto, K.R. (2005b). A Caenorhabditis elegans nutrient response system partially dependent on nuclear receptor NHR-49. Proc Natl Acad Sci U S A 102, 13496–13501, doi:10.1073/pnas.0506234102.

Vasilopoulou, C.G., Sulek, K., Brunner, A.-D., Meitei, N.S., Schweiger-Hufnagel, U., Meyer, S.W., Barsch, A., Mann, M., and Meier, F. (2020). Trapped ion mobility spectrometry and PASEF enable in-depth lipidomics from minimal sample amounts. Nat Commun 11, 331, doi:10.1038/s41467-019-14044-x.

Wanders, R.J.A., Waterham, H.R., and Ferdinandusse, S. (2015). Metabolic Interplay between Peroxisomes and Other Subcellular Organelles Including Mitochondria and the Endoplasmic Reticulum. Front Cell Dev Biol 3, 83, doi:10.3389/fcell.2015.00083.

Wang, K., Zhang, T., Dong, Q., Nice, E.C., Huang, C., and Wei, Y. (2013). Redox homeostasis: the linchpin in stem cell self-renewal and differentiation. Cell Death Dis 4, e537, doi:10.1038/cddis.2013.50.

Witting, M., Maier, T.V., Garvis, S., and Schmitt-Kopplin, P. (2014). Optimizing a ultrahigh pressure liquid chromatography-time of flight-mass spectrometry approach using a novel sub-2μm core-shell particle for in depth lipidomic profiling of Caenorhabditis elegans. J Chromatogr A 1359, 91–99, doi:10.1016/j.chroma.2014.07.021.

Xu, H.E., Lambert, M.H., Montana, V.G., Plunket, K.D., Moore, L.B., Collins, J.L., Oplinger, J.A., Kliewer, S.A., Gampe, R.T., McKee, D.D., et al. (2001). Structural determinants of ligand binding selectivity between the peroxisome proliferator-activated receptors. Proc Natl Acad Sci U S A 98, 13919–13924, doi:10.1073/pnas.241410198.

Xu, N., Zhang, S.O., Cole, R.A., McKinney, S.A., Guo, F., Haas, J.T., Bobba, S., Farese, R.V., Jr, and Mak, H.Y. (2012). The FATP1-DGAT2 complex facilitates lipid droplet expansion at the ER-lipid droplet interface. J. Cell Biol. 198, 895–911, doi:10.1083/jcb.201201139.

Young, S.G., and Zechner, R. (2013). Biochemistry and pathophysiology of intravascular and intracellular lipolysis. Genes Dev. 27, 459–484, doi:10.1101/gad.209296.112.

Zhang, S.O., Box, A.C., Xu, N., Le Men, J., Yu, J., Guo, F., Trimble, R., and Mak, H.Y. (2010). Genetic and dietary regulation of lipid droplet expansion in Caenorhabditis elegans. Proc. Natl. Acad. Sci. U.S.A. 107, 4640–4645, doi:10.1073/pnas.0912308107.

